# A Bipartite Glucan Synthase-Remodeler Module Organizes Branched Glucan Assembly in the Fungal Cell Wall

**DOI:** 10.64898/2026.02.18.706439

**Authors:** Alaina H. Willet, Anand Jacob, Lesley A. Turner, Abdulrahman Khalid A. Alsanad, Tuo Wang, Kathleen L. Gould

**Affiliations:** Department of Cell and Developmental Biology, Vanderbilt University School of Medicine, Nashville, TN 37240, USA; Department of Chemistry, Michigan State University, East Lansing, MI 48824, USA

**Author notes:** Correspondence to. Mail: Kathy Gould, PMB 407935, 465 21st Ave S, Nashville, TN 37240. These authors contributed equally (Alaina H. Willet, Anand Jacob).

## Abstract

The fungal cell wall is an essential extracellular matrix that underpins growth, morphogenesis, and pathogenesis. Cell wall construction requires numerous enzymes that synthesize and remodel extracellular polymers, yet the principles governing their spatial and functional organization remain unclear. In the fission yeast *Schizosaccharomyces pombe*, we identify Ghs2, a predicted glycoside hydrolase 16 (GH16) domain-containing transmembrane protein, as an obligate binding partner of the β-1,3-glucan synthase Bgs3. Ghs2 and Bgs3 co-localize at sites of polarized growth and physically associate *in vivo*. Structure-guided modeling positions the Ghs2 GH16 domain proximal to the predicted Bgs3 glucan extrusion pore, suggesting coordinated polymer synthesis and remodeling. Solid-state NMR analyses demonstrate that both Ghs2 and Bgs3 are required for the proper accumulation of branched β-1,3-glucan. Together with genetic and cell biological evidence, these findings support a model in which Bgs3 synthesizes linear β-1,3-glucan and Ghs2 subsequently introduces β-1,6-linked branch points onto the nascent polymer. More broadly, we propose that synthase-modifier pairs may act together to shape polymer architecture during cell wall assembly.

## Introduction

Cell walls are essential extracellular structures that provide mechanical strength, define cell shape, and protect cells from environmental stress. In walled organisms, including fungi, plants, and many microbes, proper cell wall construction is indispensable for growth, morphogenesis, and survival^1–3^. In fungi, the cell wall can comprise up to ∼20-30% of total cellular dry mass and is built from an interconnected network of polysaccharides and proteins that must be synthesized, modified, transported, and assembled outside the plasma membrane^4–6^. This process requires dozens of dedicated enzymes and accessory factors, many of which are essential^1,4,7,8^. Reflecting this complexity, cell wall assembly is among the most elaborate biosynthetic processes carried out by the cell, and 20% or more of fungal genomes encode proteins contributing to cell wall biogenesis, remodeling, and regulation^9,10^. How coordination of cell wall synthesis and remodeling is achieved to build a robust, yet adaptable extracellular matrix remains a fundamental unanswered question.

The fission yeast *Schizosaccharomyces pombe* offers a powerful system to dissect how polymer synthesis is coordinated with remodeling during cell wall assembly. In this organism, cell wall growth is spatially and temporally restricted during the cell cycle^11,12^, and the molecular composition and layered organization of the cell wall are well characterized^13–15^. The *S. pombe* cell wall consists primarily of α-1,3-glucan and β-1,3-glucan, which together account for most of the wall mass, along with smaller amounts of β-1,6-glucan and galactomannan^1,13,16–18^. The major β-1,3-glucan polymer is a branched structure composed of a β-1,3-glucan backbone with β-1,3-glucan side chains attached via β-1,6-glycosidic linkages, whereas β-1,6-glucan is a less abundant and highly branched polymer containing β-1,6-glucan side chains attached via β-1,3-glycosidic linkages^1,13,16–18^. These components are arranged into three distinct layers: an inner layer adjacent to the plasma membrane containing α- and β-glucans and galactomannan, a middle layer enriched in α- and β-glucans, and an outer layer composed mainly of glucans and galactomannan^12,13,19,20^. The enzymes that synthesize the linear β-1,3-glucan backbones that are required for the core structural scaffold of the cell wall have been determined. The biosynthesis of linear β-1,3-glucans is carried out by the essential integral membrane synthases Bgs1, Bgs3 and Bgs4 as well as the meiosis-specific Bgs2^21–25^. Despite this understanding of the enzymes that generate the β-1,3-glucan backbone, how branches are added to this polymer remains unknown^4,12^, as no specific enzyme has been shown to catalyze this remodeling step^26–32^.

Here we propose that Ghs2 is a long-sought catalytic factor required for adding branches to linear β-1,3-glucan backbones in *S. pombe*. Ghs2 is a putative single-pass transmembrane protein with an extracellular domain predicted to have cell wall-remodeling activity, and we find that it forms an obligate bipartite complex with the β-glucan synthase Bgs3. Structural modeling suggests that Ghs2 is positioned to directly add β-1,3-glucan branches to a nascent β-1,3-glucan chain via a β-1,6-linkage. This discovery establishes a paradigm for how fungi achieve tight coupling between glucan synthase activity and remodeling functions, with broad implications for cell wall biology and antifungal target discovery.

## Results

### Bgs3 and Ghs2 localization are mutually dependent

We investigated why cells with defective Ghs2 accumulate abnormal cell wall deposits^33^ by first determining the localization of glucan synthases in wildtype and *ghs2-2* mutant cells. At 25°C, GFP-tagged β-glucan synthases Bgs1, Bgs3, Bgs4, and RFP-tagged α-glucan synthase Ags1 localized to the cell tips and division site in both strains (Fig. 1a and Supplementary Fig. 1a)^34–36^. When shifted to the restrictive temperature of 36°C, Bgs1, Bgs4, and Ags1 remained properly localized in both wildtype and *ghs2-2* cells (Supplementary Fig. 1a). By contrast, GFP-Bgs3 localization was strongly reduced at cell tips and the division site in *ghs2-2* cells at 36°C (Fig. 1a, b)^35^. Total GFP-Bgs3 protein levels were unchanged in *ghs2-2* cells compared to wildtype as determined by immunoblotting (Supplementary Fig. 1b). These findings indicate that Ghs2 is specifically required for the proper subcellular localization of Bgs3, but not other glucan synthases.

**Fig. 1.**
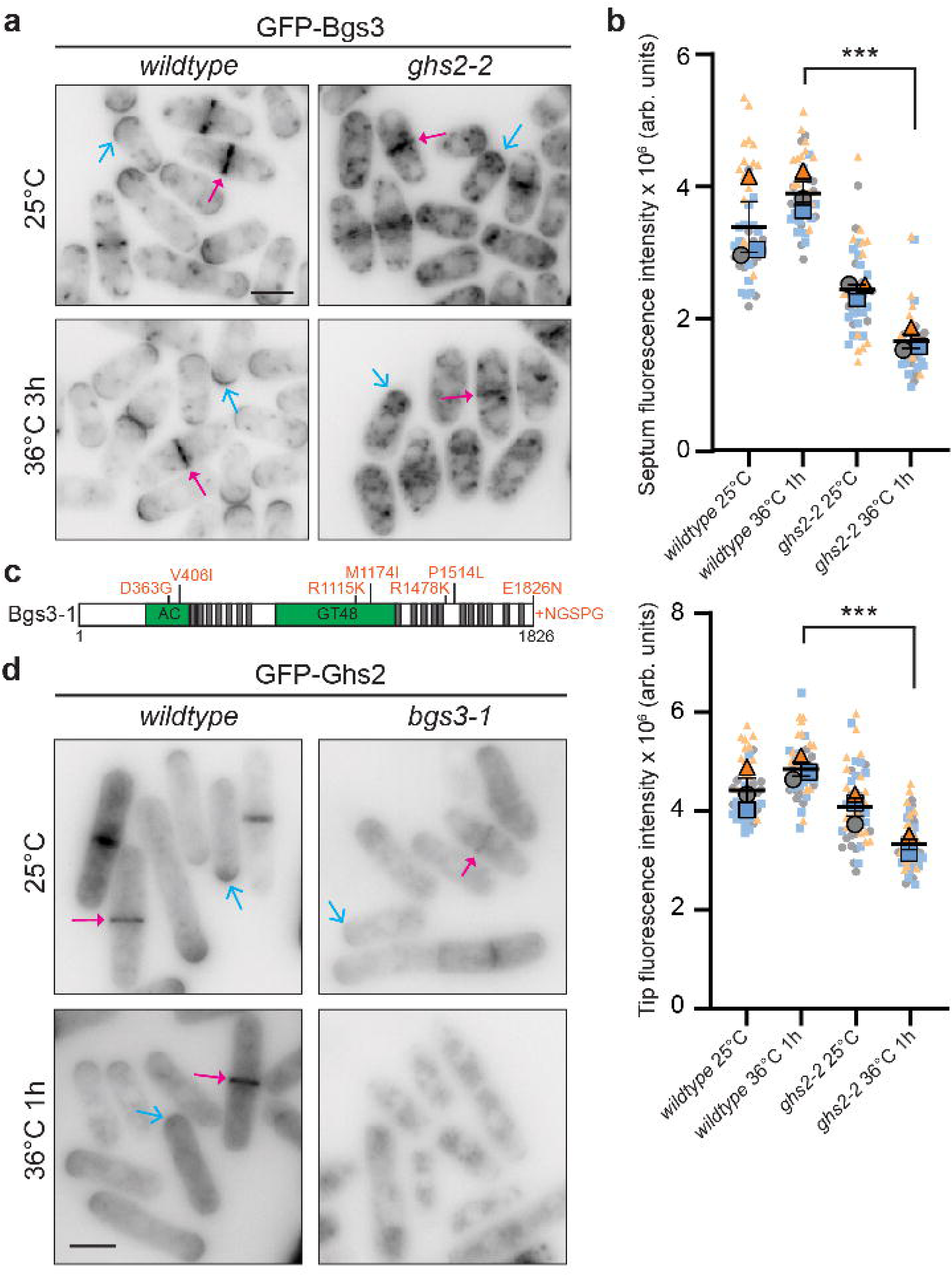
Ghs2 and Bgs3 are mutually dependent for proper localization. **a)** Representative live-cell images of GFP-Bgs3 in wildtype or *ghs2-2* cells. The strains were grown at 25°C and then shifted to 36°C for 3 hours. Images were acquired at both temperatures. Magenta arrows indicate an example cell division site localization, and cyan arrows indicate an example cell tip localization. b) Quantification of the fluorescence intensity of GFP-Bgs3 at the cell division site and cell tips in wildtype and *ghs2-2* cells at the indicated temperatures. Data are from three independent experiments. n = 45 for wildtype at both conditions at CRs and tips, n = 43 for *ghs2-2* at CRs at both conditions and n = 46 for *ghs2-2* tips at 25°C and n = 45 for *ghs2-2* tips at 36°C. Each color and symbol set (blue squares, gray circles, and orange triangles) are groups of data from the same replicate. Error bars represent mean ± SEM. Statistical analysis was performed using two-sided Welch’s *t*-test. \*\*\**P*<0.001. For the top graph, *P*=0.0004 and for the bottom graph *P*=0.001. Arbitrary units (arb. units). c) Schematic drawn to scale of the protein encoded by the *bgs3-1* allele. Mutations and additional amino acids are highlighted in orange. The accessory (AC) and glycosyltransferase 48 (GT48) domains are shown in green, and transmembrane helices are in gray. d) Representative images of wildtype and *bgs3-1* cells expressing GFP-Ghs2 from the inducible *nmt81* promoter for 24 hours prior to imaging at 25°C. The cells were then shifted to 36°C for 1 hour and imaged again. The experiment was repeated independently twice with similar results. Magenta arrows indicate GFP-Ghs2 cell division site localization and cyan arrows indicate an example GFP-Ghs2 cell tip localization. Scale bars, 5 µm.

To determine if, reciprocally, Ghs2 localization depends on Bgs3, we generated a temperature-sensitive *bgs3* allele (*bgs3-1*) (Fig. 1c). *bgs3-1* cells grew similarly to wildtype at lower temperatures but showed almost no growth at the restrictive temperature of 36°C (Supplementary Fig. 2a). In wildtype cells, GFP-Ghs2 localized to cell tips and the cell division site at both 25°C and 36°C (Fig. 1d). GFP-Ghs2 protein levels were reduced to ∼10% of wildtype levels in *bgs3-1* cells at both temperatures, as determined by immunoblotting (Supplementary Fig. 2b). This reduction could reflect decreased protein stability in the absence of functional Bgs3. Despite the reduced levels, GFP-Ghs2 still localized to cell tips and the cell division site in *bgs3-1* cells at 25°C (Fig. 1d). However, at 36°C, GFP-Ghs2 was almost entirely cytoplasmic (Fig. 1d). In contrast, GFP-Ghs2 localization was unaffected in *bgs4* (*cwg1-1*) and *bgs1* (*cps1-191*) mutants ^21,37^, and protein levels were similar to wildtype levels (Supplementary Fig. 2c). Together, these results demonstrate that Ghs2 and Bgs3 are co-dependent for proper localization to cell tips and the division site.

### Defective Bgs3 or Ghs2 function causes abnormal cell wall deposits

Like the *ghs2-2* mutant and *bgs3* knockdown cells^22,33^, *bgs3-1* cells also showed abnormal accumulation of cell wall deposits (Fig. 2a). To better understand the defects that arise in these mutants, we performed transmission electron microscopy on wildtype, *ghs2-2*, and *bgs3-1* cells grown at 25°C and shifted to 36°C for 4 hours. Wildtype cells exhibited the typical tri-lamellar cell wall architecture, with two electron-dense layers flanking a less electron-dense middle layer, as previously described^15^ (Fig. 2b, c). In contrast, both *ghs2-2* and *bgs3-1* cells had markedly similar phenotypes with thickened cell walls and large pockets of excess wall material along the lateral sides and at the tips (Fig. 2b-d). These deposits reached up to ∼600 nm in thickness, compared to the ∼120-150 nm typical thickness of wildtype cell walls (Fig. 2b-d)^19,20^. The septa of the mutant cells were also thicker than those of wildtype (Fig. 2b, e). Despite these changes, the electron-dense outer galactomannan layer remained intact (Fig. 2b, c).

**Fig. 2.**
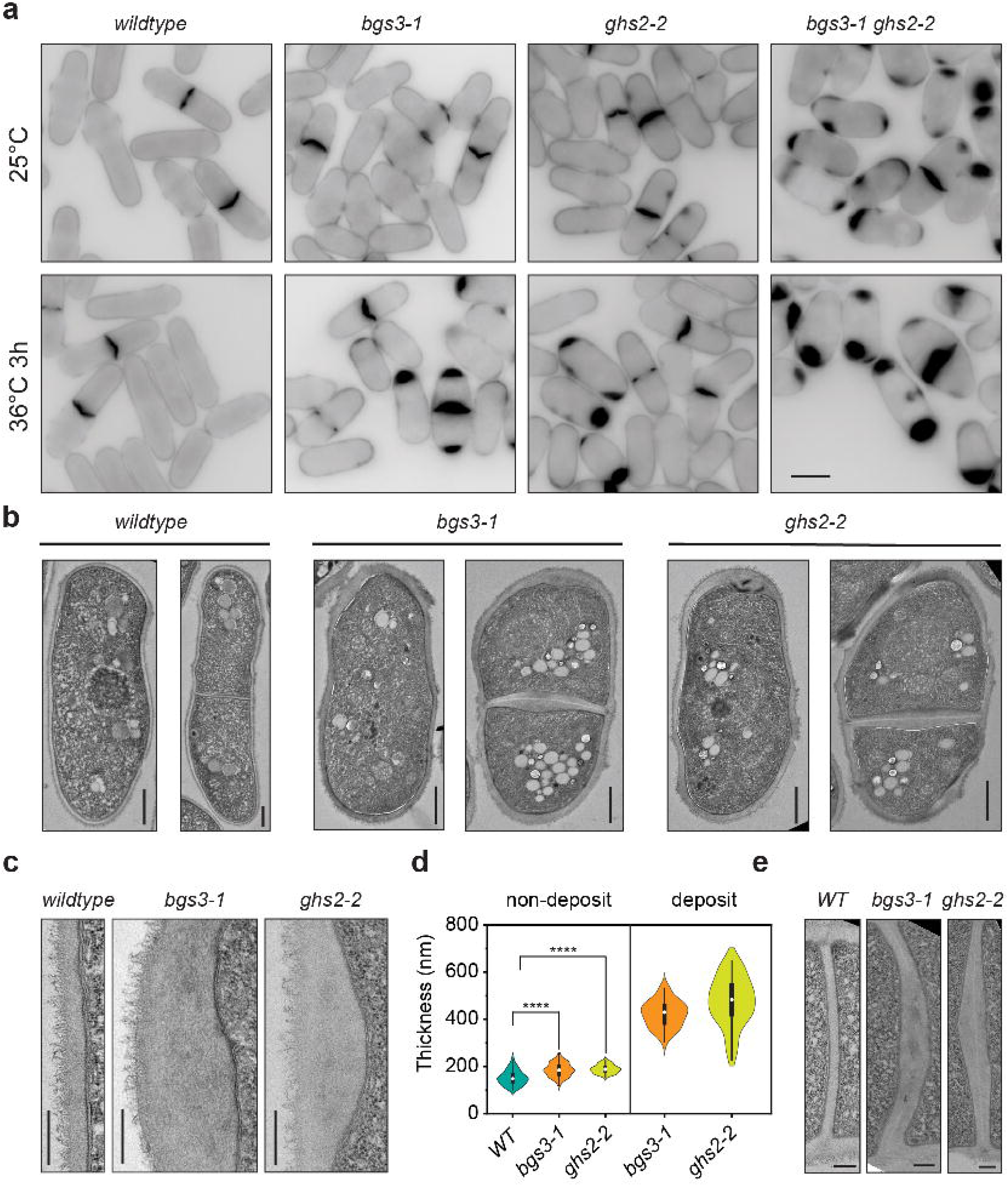
Disruption of Ghs2 or Bgs3 function results in abnormal cell wall deposit accumulation. **a)** Representative fixed-cell images of the indicated strains grown at 25°C and then shifted to 36°C for 3 hours. Cells were fixed and stained with methyl blue, a stain that binds β-glucans. b) Transmission electron microscopy (TEM) images of whole unseptated or septated wildtype, *ghs2-2* or *bgs3-1* cells grown at 25°C and shifted to 36°C for 4 hours prior to freezing the cells live. Scale bars, 100 nm. c) Zoomed-in views of the lateral cell wall of the indicated strains. Scale bars, 250 nm. d) Violin plot of cell wall thickness measured using TEM images of wildtype, *bgs3-1*, and *ghs2-2* mutant strains. Non-deposit regions were compared across all three strains (wildtype, *bgs3-1*, and *ghs2-2*), whereas deposit regions were analyzed exclusively in the two mutant strains (*bgs3-1* and *ghs2-2*). Wildtype is in green, *bgs3-1* is in orange and *ghs2-2* is in yellow. Non-deposit measurements represent 10 measurements of cell wall thickness collected from 10 individual cells (n = 100), whereas deposit measurements represent 5 measurements collected from 8 individual cells (n = 40). Black rectangles denote the interquartile range (IQR) from the 25th to 75th percentile. White circles represent the mean of the dataset, and black vertical lines correspond to 1.5 IQR. Statistical analysis for non-deposit regions was performed using single factor two-sided ANOVA between wildtype and mutant strains, with statistical significance indicated as \*\*\*\**P* < 0.0001. Between groups *P* = 3.38E-22. d) Zoomed-in views of the cell wall at fully formed septa of the indicated strains. Scale bars, 250 nm.

Given that *ghs2-2* and *bgs3-1* exhibited similar defects, it was not unexpected that combining these two hypomorphic alleles led to enhanced growth defects (Figure 2a and Supplementary Fig. 2a), with more and larger cell wall deposits than observed in either single mutant alone (Fig. 2a and Supplementary Fig. 2d, e). These findings are consistent with Bgs3 and Ghs2 functioning together to maintain normal cell wall architecture.

### A Ghs2-Bgs3 complex integrates β-1,3-glucan synthesis with extracellular remodeling

Because Ghs2 and Bgs3 were interdependent for localization, it was not surprising to find that mCherry-Bgs3 and GFP-Ghs2 co-localized at cell tips, the division site, and in internal puncta (Fig. 3a). Furthermore, consistent with Ghs2 specifically associating with Bgs3, HA_3_-Ghs2 and GFP-Bgs3 co-immunoprecipitated (Fig. 3b) whereas GFP-Ghs2 and RFP-Bgs4 did not (Supplementary Fig. 3a).

**Fig. 3.**
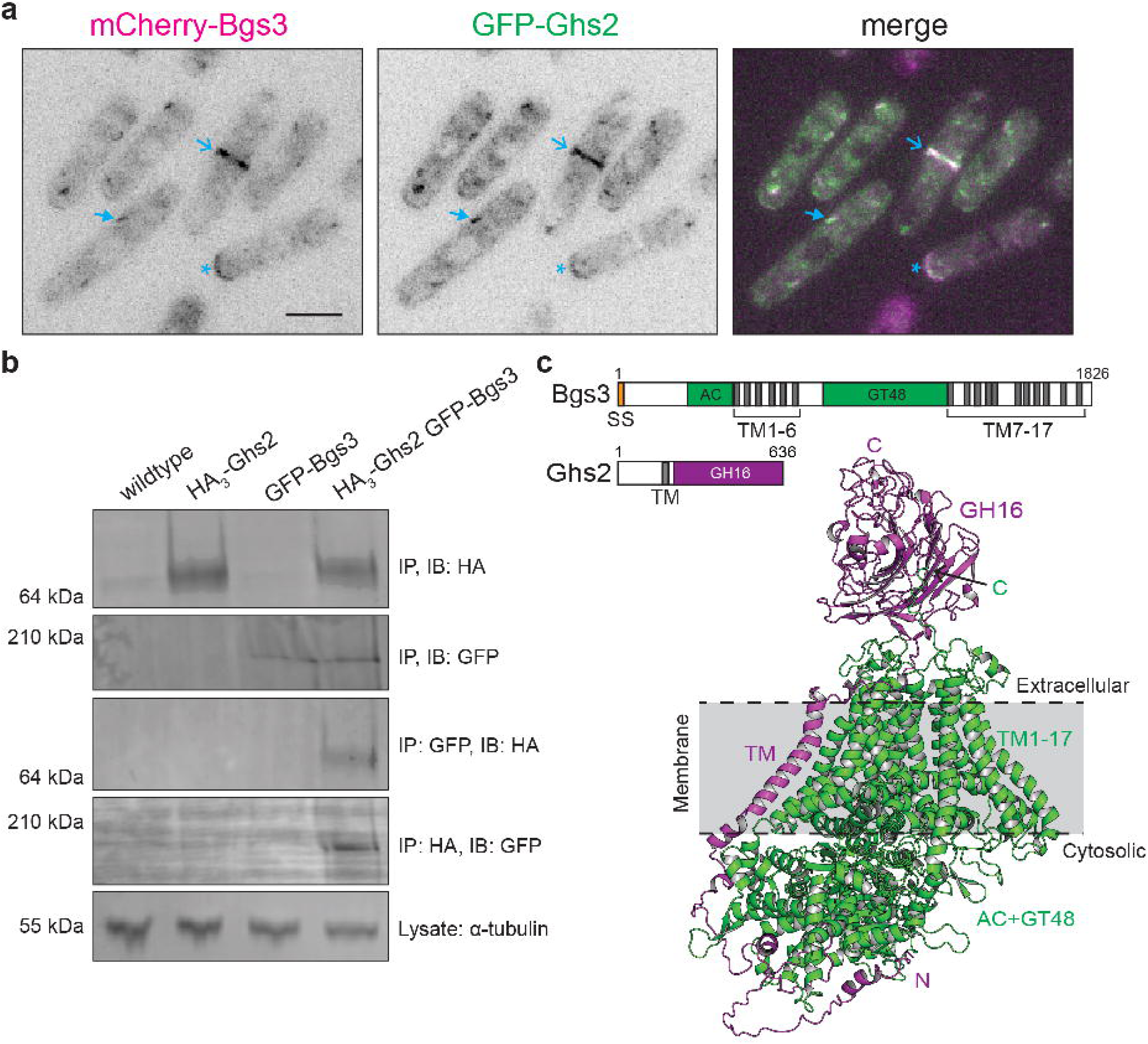
Bgs3-Ghs2 associate in cells. **a)** Representative live-cell imaging of mCherry-Bgs3 cells expressing *nmt81-GFP-ghs2* for 24 hours prior to imaging at 25°C. The experiment was repeated independently twice with similar results. Open arrows denote co-localization at the cell division site, asterisks indicate co-localization at the cell tip and closed arrows mark co-localization at internal puncta. Scale bar, 5 µm. b) Anti-HA and anti-GFP immunoprecipitations from the indicated strains. Samples were blotted with anti-GFP or anti-HA antibodies. Lysate samples were blotted for α-tubulin (bottom). The experiment was repeated independently twice with similar results. c) Top, schematics, drawn to scale, of Bgs3 and Ghs2. Transmembrane (TM) helices are in gray, the accessory domain (AC) and glycosyl transferase 48 (GT48) domains are in green, and the glycoside hydrolase 16 (GH16) domain is in purple and the signal sequence (SS) is in orange. Bottom, AlphaFold3-generated model of Bgs3-Ghs2 complex. Bgs3 is in green and Ghs2 is in purple.

To probe the Ghs2-Bgs3 interaction in more detail, we used AlphaFold3 (AF3)^38^. AF3 modeling suggests multiple direct binding interfaces including one involving the unstructured N-terminus of Ghs2 and the intracellular catalytic domain of Bgs3, a second involving contacts between the Ghs2 transmembrane (TM) helix and TM helices 6 and 8 of Bgs3, and a third involving the Ghs2 GH16 domain and extracellular loops between TM helices of Bgs3 (Fig. 3c and Supplementary Fig. 3b). Strikingly, AF3 positioned the GH16 domain of Ghs2 on the extracellular side of the membrane, proximal to the predicted β-1,3-glucan extrusion pore of Bgs3 (Fig. 3c)^39,40^.

Intriguingly, the Ghs2 TM domain shares sequence similarity with the TM domain of Sbg1^41^ (Supplementary Fig. 3c), which associates with the β-1,3-glucan synthase Bgs1 and is required for Bgs1’s proper localization and function at sites of polarized growth^42,43^. Given this similarity, we asked whether AF3 would predict a comparable mode of association between Sbg1 and Bgs1 as observed for Ghs2 and Bgs3. Indeed, AF3 predicted an analogous architecture of the complex (Supplementary Fig. 3d, e).

Unlike Sbg1, Ghs2 contains an extracellular domain with predicted catalytic function, the GH16 domain. We generated a series of *ghs2* mutants to separate the requirements for Bgs3 binding from those required for GH16 enzymatic activity. First, to test the functional role of the Ghs2 TM helix, we replaced the native Ghs2 TM helix with the homologous TM helix from Sbg1 or the unrelated TM helix from the plasma membrane protein Wsc1 (Fig. 4a). The TM helices from these three proteins are all predicted to be 23 residues in length^44^. Strikingly, substitution with the Sbg1 TM domain supported growth in *ghs2-2* cells at the non-permissive temperature, however substitution with the Wsc1 TM helix did not (Fig. 4a). This result reinforces the sequence relatedness of the Ghs2 and Sbg1 TM domains.

**Fig. 4.**
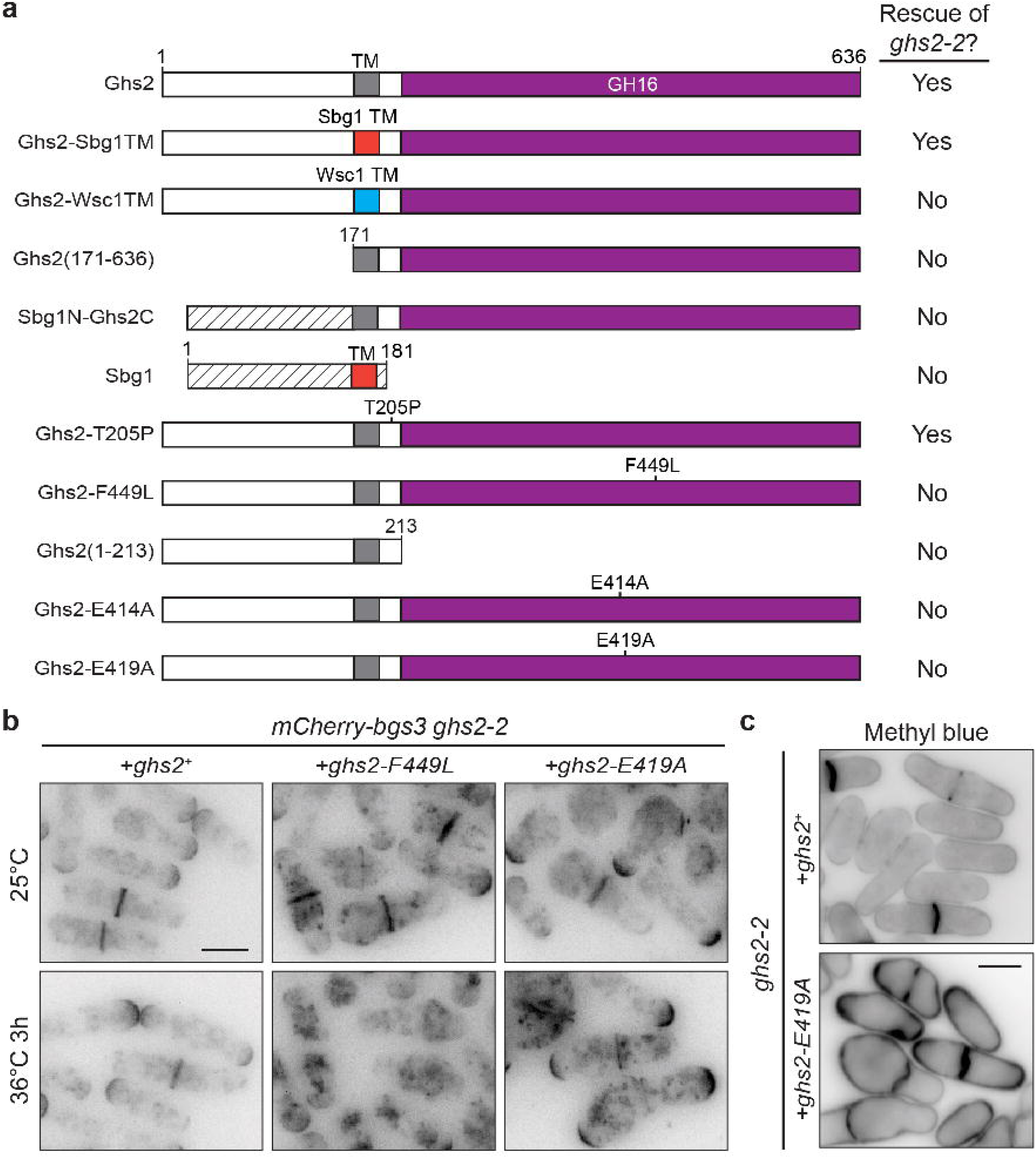
Ghs2 TM helix and GH16 domain are essential for function. **a)** Schematics, drawn to scale, of the indicated proteins expressed from the *nmt1* promoter. The Ghs2 glycoside hydrolase 16 (GH16) domain is in magenta, the Ghs2 transmembrane (TM) helix is in gray, the Sbg1 TM helix is in orange and the Wsc1 TM helix is in cyan. The Sbg1 protein sections, other than the TM helix, are indicated with black stripes. The plasmids containing the genes encoding the indicated proteins were transformed into *ghs2-2* cells, grown at 25°C and then struck out at 36°C to test for rescue in the presence of thiamine. b) Live-cell imaging of *mCherry-bgs3 ghs2-2* cells expressing an empty vector or the indicated *ghs2* constructs. Cells were grown at 25°C and shifted to 36°C for 3 hours and imaged at both time points. The experiment was repeated independently twice with similar results. c) Representative fixed-cell images of the indicated *ghs2-2* cells expressing *ghs2^+^* or *ghs2-E419A* and grown at 25°C. Cells were fixed and stained with methyl blue. The experiment was repeated independently twice with similar results. Scale bars, 5 µm.

Further examination of the AF3-predicted complex suggested that the conserved Ghs2 TM helix residues L177, L184 and F185 mediated binding to Bgs3 residues W670 and L673 within helix 6 and residues I1224, L1341 and I1338 within helix 8 (Supplementary Fig. 4a, b). To test this interaction model, we mutated these residues to alanine and tested their ability to rescue growth of their respective temperature-sensitive allele. We found that neither *ghs2-3A* (L177A, L184A, F185A) nor *bgs3-5A* (W670A, L673A, I1334A, L1341A, I1338A) complemented growth of *ghs2-1* or *bgs3-1*, respectively (Supplementary Fig. 4c).

We then asked whether the Ghs2 N-terminal cytosolic domain was interchangeable with the N-terminus of Sbg1. We found that the N-terminus is required for Ghs2 function and replacing it with that of Sbg1 failed to complement growth of *ghs2-2* cells (Fig. 4a).

Overexpression of full-length Sbg1 likewise did not rescue the *ghs2-2* mutant suggesting that unlike the TM helices, the N-termini of these two proteins have specialized functions (Fig. 4a), likely dictating which glucan synthase they interact with.

We next examined the role of the predicted Ghs2 extracellular region. We started by testing the individual contributions of the two mutations in the *ghs2-2* allele, T205P located between the TM helix and the GH16 domain, and F449L located within the GH16 domain^33^. *ghs2-T205P* supported growth in *ghs2-2* cells at the non-permissive temperature, whereas *ghs2-F449L* did not (Fig. 4a), indicating that F449L is the causative mutation of the *ghs2-2* conditional lethal phenotype. F449 is predicted to be a buried residue within the GH16 fold^45,46^, and therefore this substitution likely destabilizes the domain and the protein. In accord with the GH16 domain being important for Ghs2 function, we found that a construct comprising only the N-terminus and TM domain, similar to the architecture of Sbg1, did not rescue *ghs2-2* (Fig. 4a).

To specifically disrupt GH16 catalytic activity without disturbing domain structure, we introduced substitutions (E414A or E419A) within the conserved catalytic ^414^EXDXXE^419^ motif^47^ previously shown to inactivate but not destabilize a bacterial GH16 enzyme (Fig. 4a)^48^. Neither mutant rescued growth of *ghs2-2* cells (Fig. 4a). To assess how these mutations affect Bgs3, we examined mCherry-Bgs3 localization in *ghs2-2* cells expressing *ghs2^+^*, *ghs2-F449L*, or *ghs2-E419A* (Fig. 4b). At 36°C, mCherry-Bgs3 was diminished at cell tips and division sites in *ghs2-F449L* cells compared to wildtype, consistent with what we had observed in the *ghs2-2* mutant (Fig. 4a, b and Fig. 1a). In contrast, mCherry-Bgs3 localized robustly to cell tips and the cell division site in cells expressing *ghs2-E419A* at 36°C (Fig. 4a, b). These observations support the interpretation that E419A preserves the stability of the Ghs2-Bgs3 complex but abolishes Ghs2 GH16 catalytic activity. In support of this interpretation, expression of *ghs2-E419A* in *ghs2-2* cells at the permissive temperature caused severe cell wall defects and accumulation of β-glucans (Fig. 4c). This indicates that loss of GH16 catalytic activity, while leaving complex formation with Bgs3 intact, is sufficient to drive cell wall dysfunction. The strong deleterious phenotype also suggests that *ghs2-E419A* behaves as a dominant-negative allele, underscoring that both the formation of the Ghs2-Bgs3 complex and Ghs2 enzymatic activity are indispensable for proper cell wall assembly.

### Ghs2 and Bgs3 are required for cell wall branched β-1,3-glucan linkages

Bgs3 is thought to contribute to cell wall assembly by synthesizing linear β-1,3-glucan chains^22^. In contrast, the biochemical role of Ghs2 is unclear although GH16 domains typically hydrolyze or transglycosylate β-glucans at various linkage sites^47^. We employed solid-state NMR spectroscopy^13,49–51^ to analyze the composition of intact, natively hydrated cell walls of wildtype, *bgs3-1*, *ghs2-2,* and *bgs3-1 ghs2-2* cells to more precisely define the roles of Ghs2 and Bgs3.

The fungal cell wall is composed of regions with distinct physical properties, including a rigid phase and a mobile phase^1,5,10,52^. In *S. pombe*, the rigid phase consists of densely packed, stiff molecules that occupy the central core of the cell wall, whereas the mobile phase is composed of more flexible molecules enriched in the membrane-proximal and distal regions of the wall^53–57^. By tailoring solid-state NMR methodologies, rigid and mobile components of the fungal cell wall can be selectively resolved based on their intrinsic molecular dynamics. This capability enables direct, noninvasive analysis of intact cell walls and has established solid-state NMR as a central tool for elucidating fungal cell wall composition and material properties. Its application has driven major advances in understanding cell wall architecture across diverse fungal taxa, including *Aspergillus*^52,57–61^, *Cryptococcus*^62,63^, *Candida*^64,65^, *Schizophyllum*^66–68^, and members of the *Rhizopus* and *Mucor* species^69^. Recent studies defined the *S. pombe* rigid cell wall phase as being primarily composed of β-1,3-glucan and α-1,3-glucan with a minor β-1,6-glucan population, and the mobile phase contains these polymers as well as mannan polymers containing α-1,2- and α-1,6-linked mannose units^13,70^. Initial 1D solid-state NMR spectra of intact cells revealed distinct alterations in the cell wall composition of both *ghs2-2* and *bgs3-1* single mutants compared to wildtype (Supplementary Fig. 5-7). The signals corresponding to the major β-1,3-glucan population and the minor β-1,6-glucan population were reduced within the mobile domain, accompanied by a decrease in protein resonances from the rigid core (Supplementary Fig. 5, 6). The compositional changes were more pronounced in the *bgs3-1 ghs2-2* double mutant, which exhibited an extensive loss of β-1,3- and β-1,6-glucans, near-complete depletion of mobile carbohydrate components (Supplementary Fig. 6c), and disappearance of protein peaks from the rigid phase (Supplementary Fig. 5c).

To obtain higher-resolution insight into these alterations, 2D solid-state NMR analyses were performed on *bgs3-1, ghs2-2* and *bgs3-1 ghs2-2* cells after a 4 h shift to the non-permissive temperature (Fig. 5a-c, Supplementary Fig. 8). Within the rigid cell wall core, all three mutants, selectively probed via dipolar-coupling-based methods, lacked detectable β-1,6-glucan and displayed a new polymorphic population of β-1,3-glucan (B^b^), which was absent from wildtype cells (Fig. 5c). Within the mobile domains, selectively detected using their rapid ^13^C spin-lattice relaxation, the β-1,3,6-glucopyranose containing branched glucan (B^Br^) of *bgs3-1* and *ghs2-2* was reduced by ∼58% and ∼17%, respectively and, β-1,6-glucan was reduced by ∼45% and ∼30%, respectively (Fig. 5c, d). The β-1,3,6-glucopyranose-containing branched glucan (B^Br^) represents glucan residues that are substituted at both the 3- and 6-positions and therefore may arise from either branched β-1,3-glucan or branched β-1,6-glucan polymers (Fig. 5d). In addition, *bgs3-1*, but not *ghs2-2,* also had an ∼8-fold increase in β-1,3-glucan (Fig. 5c).

**Fig. 5.**
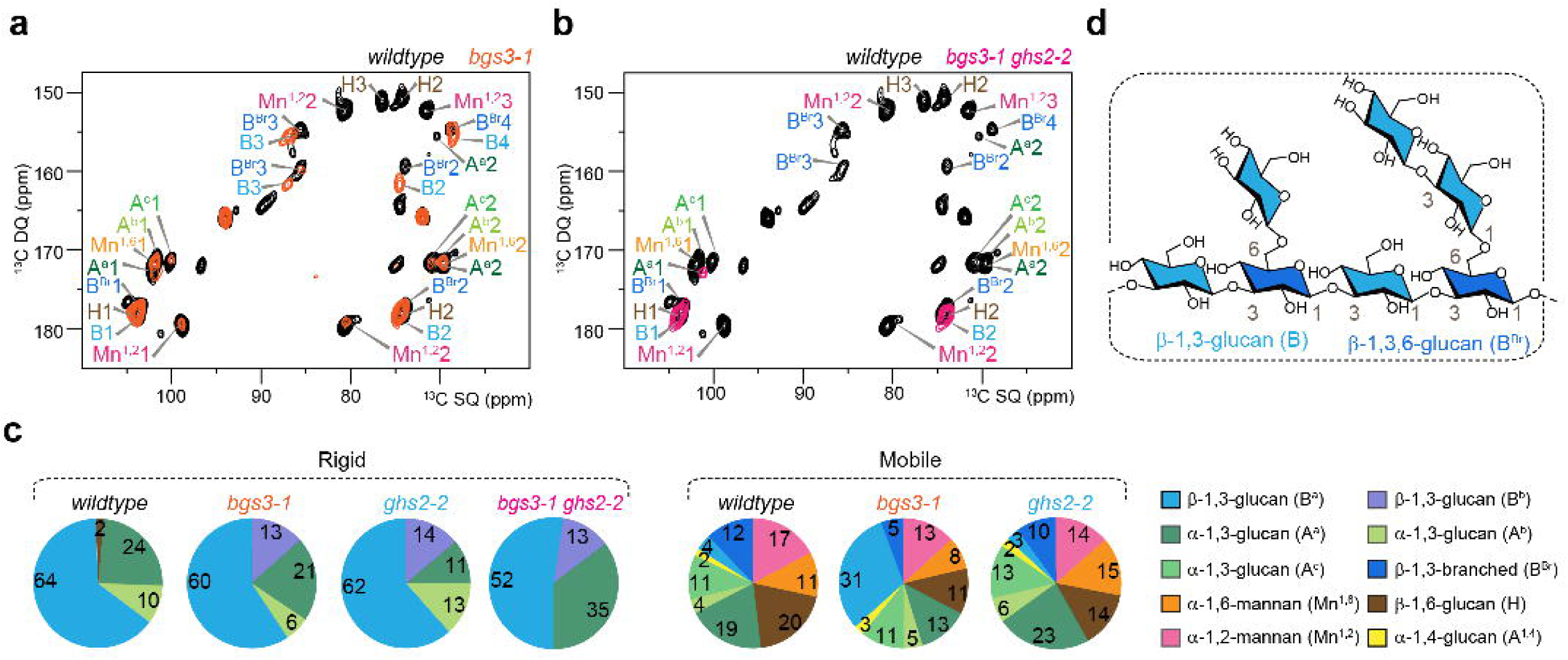
Comparative analysis of cell wall composition in wildtype, *bgs3-1, ghs2-2* and *bgs3-1 ghs2-2* mutant cell walls. 2D ^13^C DP refocused J-INADEQUATE spectra detecting the mobile polysaccharides of wildtype (black) compared with a) *bgs3-1* (orange) and b) *bgs3-1 ghs2-2* (pink). The spectra revealed extensive disruption of mobile polysaccharides in the *bgs3-1 ghs2-2* strain. c) Relative molar composition of rigid and mobile polysaccharides determined from integrated cross-peak volumes in the 2D ^13^C-^13^C CORD and 2D ^13^C DP refocused J-INADEQUATE spectra respectively and are expressed as molar percentages (%). d) A simplified architecture of β-1,6-branched β-1,3-glucan that is present in the *S. pombe* cell wall.

The mobile fraction of the double mutant was dramatically remodeled compared to wildtype. The absence of 2D ^13^C DP-INADEQUATE signals in the double mutant indicates that the flexible fraction of the cell wall is largely lost (Fig. 5b). This reflects a substantial reduction or immobilization of β-1,3- and β-1,6-glucans and the loss of α-1,2- and α-1,6-linked mannose residues, which were readily detected in wildtype. With the outer galactomannan layer disrupted there was insufficient mobile material to produce detectable through-bond correlations and thus calculation of relative molar composition of mobile polysaccharides (Supplementary Fig. 6). Together, these findings identify the Ghs2-Bgs3 complex as the key determinant of branched β-glucan and β-1,6-glucan content and indicate that disruption of their partnership broadly reorganizes the architecture of the cell wall.

### D75-4590 treatment produces a Ghs2 loss-of-function-like phenotype in S. pombe

Several small-molecule inhibitors are proposed to directly target the catalytic GH16 domain of Ghs2 homologs, like *S. cerevisiae* Kre6^71,72^. These compounds include the steroidal alkaloid jervine and related compounds as well as D75-4590 and its derivatives^71,73–75^. Kre6-related proteins are the expected target of these inhibitors based on genetic evidence: a phenylalanine-to-isoleucine substitution in the Kre6 GH16 domain (F552I) confers resistance to both D75-4590 and jervine in *S. cerevisiae*^71,72^. D75-4590 was originally identified in a screen for compounds that cause loss of cell wall-associated proteins covalently crosslinked to β-1,6-glucan^71^, and loss of Kre6 reduces β-1,6-glucan levels across multiple fungal species^32,76–79^.

Consequently, these compounds have been interpreted as inhibitors of β-1,6-glucan biogenesis/synthesis however, there is no direct biochemical evidence defining the specific enzymatic activity inhibited by these compounds.

*S. pombe* Ghs2 and *S. cerevisiae* Kre6 are ∼46% identical, share conserved domain architectures, and contain GH16 domains that superimpose closely (root-mean-square deviation of Cα atom positions = 0.42 Å over 331 residues) (Supplementary Fig. 9a, b)^45,46,80^. To test whether D75-4590 exhibits antifungal activity against *S. pombe*, wildtype cells were treated with a range of drug concentrations (0.25-4 mM) and we determined the MIC-2 (defined as the concentration that reduces cell growth by 50% after 24 h) to be 4 mM (Table 1). We report MIC-2 rather than MIC due to the limited solubility of D75-4590 in aqueous solution, consistent with prior studies, and the relatively high concentration needed to observe growth inhibition^74^. Imaging of wildtype cells treated with a range of D75-4590 concentrations (0.125-4 mM) revealed a dose-dependent accumulation of aberrant cell wall material at the growing cell tips and cell division site (Fig. 6a). This phenotype closely resembled the temperature-dependent defects observed in *ghs2-2* cells (Fig. 2a). In accord, jervine-treated *S. cerevisiae* cells also accumulated abnormal β-glucans in the daughter bud^72,75^.

**Fig. 6.**
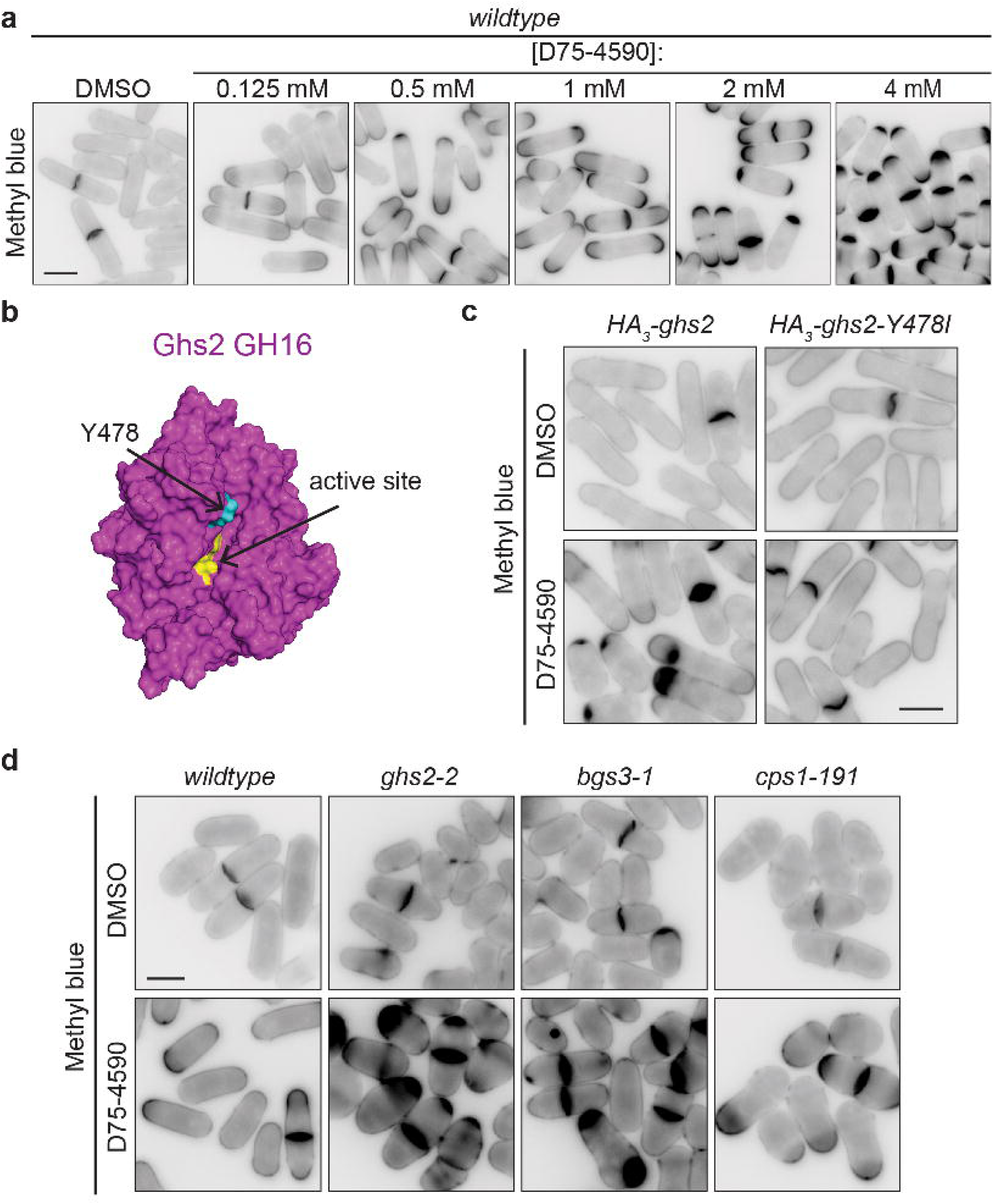
Genetic and pharmacological evidence implicates Ghs2 as a target of the small-molecule inhibitor D75-4590. **a)** Representative fixed-cell images of wildtype cells grown at 29°C in EMM and treated with DMSO or the indicated concentration of D75-4590 for 3 hours prior to fixing and staining with methyl blue. The experiment was repeated independently twice with similar results. b) AlphaFold3-generated structure of the Ghs2 GH16 domain in magenta. The conserved catalytic ^414^EXDXXE^419^ motif is in yellow and residue Y478 is in cyan. c) Representative fixed-cell images of *HA_3_-ghs2^+^* or *HA_3_-ghs2-Y478I* cells grown in EMM at 27°C treated with DMSO or 1 mM D75-4590 for 3 hours. Cells were then fixed and stained with methyl blue. The experiment was repeated independently three times with similar results. d) Representative fixed-cell images of wildtype, *ghs2-2, bgs3-1* or *cps1-191* cells grown in YE at 27°C treated with DMSO or 1 mM D75-4590 for 3 hours. Cells were fixed and stained with methyl blue. The experiment was repeated independently twice with similar results. Scale bars, 5 µm.

**Table 1.** In vivo susceptibility to D75-4590. The minimal inhibitory concentration (MIC-2) was determined as the concentration of D75-4590 where there was ∼50% of growth compared to control continuing only DMSO for each strain. Cells for each strain were diluted to a density of 3×10^6^ cells/mL in minimal medium containing appropriate supplements at 29°C. Each condition contained 0, 0.125, 0.25, 0.5, and 1 mM of D75-4590. The wildtype sample was also treated with 2 and 4 mM of D75-4590. The indicated values are the average from three independent experiments. The absorbance data was represented as: −(0-4.9% turbidity), + (5-9.9% turbidity), ++ (10-24.9% turbidity), +++ (25-49.9% turbidity), ++++ (50-74.9% turbidity), and +++++ (75-100% turbidity). Not applicable (N/A).

| Strains | [D75-4590] (mM) |  |  |  |  |  |  |
| --- | --- | --- | --- | --- | --- | --- | --- |
|  | 0 | 0.125 | .25 | .5 | 1 | 2 | 4 |
| *wildtype* | +++++ | +++++ | +++++ | +++++ | ++++ | ++++ | +++ |
| *HA<sub>3</sub>-ghs2* | +++++ | ++++ | +++ | +++ | ++ | N/A | N/A |
| *HA<sub>3</sub>-ghs2-Y478I* | +++++ | +++++ | +++++ | +++++ | +++ | N/A | N/A |

*S. cerevisiae* Kre6 F552, the residue that when mutated to isoleucine confers resistance to D75-4590 and jervine, is analogous to *S. pombe* Ghs2 Y478 and these residues are predicted to occupy structurally equivalent positions at the entrance to the GH16 domain substrate-binding cleft (Fig. 6b and Supplementary Fig. 9b)^45,46^. To test whether Ghs2 Y478 similarly contributes to *S. pombe* inhibitor sensitivity, we introduced the analogous Y478I substitution into Ghs2. The control *HA_3_-ghs2* strain had an MIC-2 of 0.25 mM, while *HA_3_-ghs2-Y478I* cells had a much higher MIC-2 of 1 mM (Table 1). In corroboration, both strains grew similarly on DMSO-containing plates, but *HA_3_-ghs2-Y478I* grew better than *HA_3_-ghs2* on plates containing 0.25 mM D75-4590 (Supplementary Fig. 10a). Upon D75-4590 treatment, *∼*76% (n = 329) of *HA_3_-ghs2* cells accumulated abnormal cell wall material, while only ∼6% (n = 311) *HA_3_-ghs2-Y478I* cells did (Fig. 6c). Finally, we tested whether *bgs3-1* and *ghs2-2* mutants were hypersensitive to D75-4590. Both D75-4590-treated *ghs2-2* and *bgs3-1,* but not the temperature-sensitive allele of *bgs1* (*cps1-191*), exhibited a higher percentage of cells that accumulated abnormal cell wall material and formed larger cell wall deposits than wildtype cells (Fig. 6d and Supplementary Fig. 10b, c). Together, these data indicate that Y478 contributes to D75-4590 sensitivity, supporting a conserved inhibitor-binding site within the GH16 domain.

## Discussion

Here, we show that Ghs2 partners physically and functionally with the β-1,3-glucan synthase Bgs3, and propose that it functions as a remodeling enzyme which introduces branches onto linear β-1,3-glucan chains in *S. pombe*. Ghs2-related proteins exist in other fungi, but they have not been demonstrated to directly couple to any cell wall synthase enzyme^4,81^.

Ghs2 and Bgs3 are strictly interdependent for localization, co-purify, and structural modeling suggests that they form a complex with the Ghs2 GH16 domain ideally positioned to introduce β-1,3-glucan branches onto newly synthesized β-1,3-glucan chains as they are extruded into the periplasmic space by Bgs3 (Fig. 7a). Our findings establish a paradigm for how fungi can achieve tight coupling between glucan synthase activity and remodeling functions, with broad implications for cell wall biology and antifungal target discovery.

**Figure 7.**
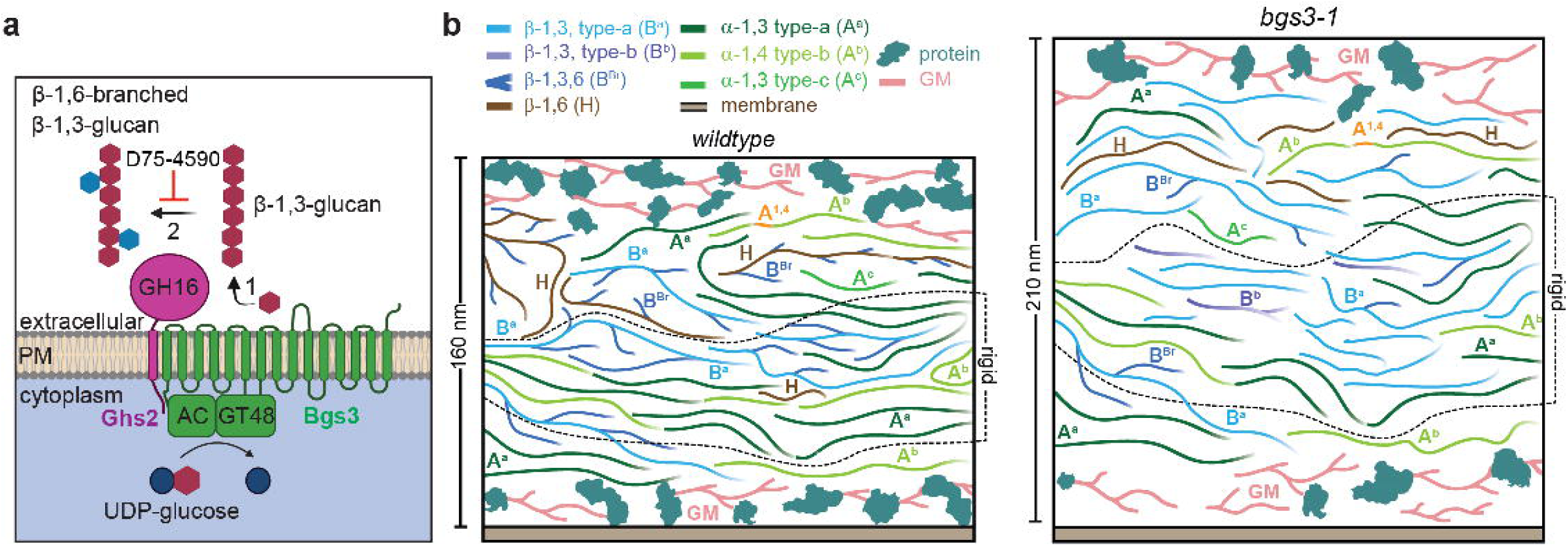
Proposed model for the function of the Ghs2-Bgs3 complex. **a)** We propose a model where β-1,3-glucan (red) is synthesized and extruded by the β-glucan synthase Bgs3 (green). Then Ghs2 (magenta), which is bound to Bgs3, is positioned to modify nascent β-1,3-glucan by adding β-1,6-glucan-linked branches (blue) of β-1,3-glucan chains. We also propose that the D75-4590 small molecule inhibitor inhibits this branching activity of Ghs2 and related proteins. Created in BioRender. Willet, A. (2026) https://BioRender.com/x9ntpno. b) Illustrative cartoons show simplified *S. pombe* cell wall architectures in wildtype and *bgs3-1* cells. The legend identifies four β-glucan species, three α-glucan species, galactomannan (GM), membrane, and cell wall proteins. In each panel, the region between the dashed lines denotes the rigid wall core, whereas the flanking regions represent the mobile fractions. Plots summarize the molecular composition of the rigid and mobile domains (not drawn to scale). The schematics integrate measurements of wall thickness from imaging with compositional and structural information from solid-state NMR, together with prior biochemical data and established models of the *S. pombe* cell wall.

Prior studies have suggested that Ghs2/Kre6-related enzymes may function in the ER-Golgi system, whereas others support a role at the cell surface^19,77,82,83^. Our data support a model in which Ghs2 traffics through the secretory pathway but functions at the plasma membrane together with Bgs3. Consistent with this model, immuno-electron microscopy images of *S. pombe* have shown that branched β-1,3-glucan chains extend outward from the plasma membrane into the wall matrix^19^. Our solid-state NMR analyses further demonstrated that loss of either Ghs2 or Bgs3 led to reduced β-glucan branching (Fig. 7b), suggesting that this synthase-modifier pair is the source of branched β-1,3-glucan within the cell wall.

Our modeling and genetic evidence further support that the Ghs2 GH16 domain performs a branching reaction. Notably, the Y478I substitution in Ghs2 confers resistance to an antifungal drug that targets Ghs2/Kre6-related proteins. Structural models place Y478 at the entrance to the catalytic cleft, where it helps shape the channel through which substrates access the active site. In this position, the aromatic side chain of Y478 could contribute to ligand recognition by stabilizing bound molecules at the mouth of the pocket. Replacing tyrosine with isoleucine maintains a hydrophobic residue at this position, which may allow normal substrate entry, but removes the aromatic ring that an inhibitor might rely on for binding. This change would be expected to reduce inhibitor interaction without disrupting catalytic function. Together, the modeling provides an explanation for the resistance phenotype: the mutation alters the architecture of the active-site entrance in a way that weakens inhibitor binding while preserving enzymatic activity. Further, we propose that D75-4590 inhibits the branching activity of Ghs2/Kre6 homologs and the loss of β-1,6-glucan polymers may be a downstream consequence of inhibiting Ghs2/Kre6 activity.

Disruption of either Ghs2 or Bgs3 results in a reduction of β-1,6-glucan in both rigid and mobile cell-wall fractions. While the β-1,6-glucan signal in the mobile fraction represents a highly branched polymer, we detected a small percentage (∼2%) of linear β-1,6-glucan in the rigid core of the cell wall. The reduction in cell wall proteins and thicker cell walls observed in the *bgs3* and *ghs2* mutants are consistent with downstream consequences of disrupted β-1,6-glucan and with the known crosslinking and scaffolding role of β-1,6-glucan in diverse fungi^4,5,26,84^. The loss of both β-1,6-glucan populations is not readily explained by the loss of branched β-1,3-glucan alone (Fig. 7b). This may suggest that branched β-1,3-glucan is required for the proper incorporation, organization, or stability of the β-1,6-glucan content within the wall matrix and that branch points introduced by Ghs2 may generate structural features that facilitate the retention or crosslinking of these components by additional enzymes.

A key outstanding question however is how the Ghs2 GH16 domain would generate the β-1,6-glycosidic linkages that attach β-1,3-glucan side chains to the β-1,3-glucan backbone. Our data point to a transglycosylation mechanism in which Ghs2 transfers one or more glucosyl residues from a β-1,3-glucan donor onto the O6 position of nascent Bgs3-extruded β-1,3-glucan chain (Fig. 6d). Classic linkage analysis from *S. cerevisiae* suggests that most of these branches consist of very short stubs, typically single residues, with occasional two- or three-residue extensions^85^.

Our data indicate that the TM domains of Ghs2 and Sbg1 are interchangeable units for glucan synthase association. The conservation of Ghs2/Sbg1-like TM domains across fungi (Supplementary Fig. 3b) raises the possibility that proteins containing them also bind cell wall synthases in other species. Sbg1 is critical for the trafficking of Bgs1 to the plasma membrane^42,43^. It is likely that Ghs2 and Bgs3 are also trafficked together because they co-localize at the plasma membrane and in intracellular puncta and are reciprocally required for the other’s plasma membrane localization. Therefore, interaction with Sbg1 or Ghs2 might provide a trafficking signal for the glucan synthases and/or confer stability. The N-terminal intracellular regions of Sbg1 and Ghs2 appear to encode glucan synthase binding preference, and further studies are needed to identify the precise specificity cues. No analogous Sbg1/Ghs2 partners have been identified for the other *S. pombe* β-glucan synthases, Bgs2 and Bgs4. Interestingly, Bgs4 associates with Smi1, an intracellular protein lacking both a transmembrane segment and any predicted catalytic domain^86^, suggesting that distinct classes of auxiliary factors may regulate different glucan synthases and support a division of labor in cell wall construction: Bgs1, together with Sbg1, plays a central role in linear β-1,3-glucan synthesis for primary septum formation and are also important for polarized growth at the cell tips, Bgs4 contributes to β-1,3-glucan synthesis in both the septum and lateral cell wall, and Bgs3, together with Ghs2, promotes the formation of the branched β-1,3-glucan network at the septum and along the lateral cell wall. This specificity would explain why over-expression of other β-glucan synthases cannot compensate for Bgs3 loss, and reciprocally, Bgs3 cannot substitute for other synthases^12,21,22,36,87^.

Beyond *S. pombe*, comparative genomics highlight that certain oomycetes encode a multidomain protein in which a β-1,3-glucan synthase domain is fused to a GH16 domain within a single polypeptide^88,89^. Phylogenomic analyses indicate that this architecture is restricted to oomycetes and absent from fungi, plants, and animals, suggesting it represents a lineage-specific innovation rather than an ancestral state. Thus, fungi and oomycetes appear to have convergently evolved distinct strategies to achieve the same outcome of tight coupling between β-glucan synthesis and remodeling. A similar principle is seen in fungi with Ags1, which integrates α-glucan synthase and remodeling activities^13,90^. Together, these examples illustrate multiple organizational strategies that ensure coordinated synthesis and remodeling, highlighting how fungi evolved modular pathways for robust extracellular matrix construction, with direct implications for antifungal target discovery.

## Methods

### Yeast strains and media

*S. pombe* strains (Supplementary Table 1) were grown in yeast extract (YE) medium or minimal medium (EMM) with appropriate supplements^91^. Transformations were performed by the lithium acetate method^92^. G418 (Geneticin, 100 µg/mL, Thermo Fisher Scientific; cat. # 11811031) was used for selection of strains containing appropriate tags. Strain construction and tetrad analyses were accomplished through standard methods^93^. Unless otherwise noted, cells used for live cell imaging were grown at 25°C in YE medium.

Multiple strategies were employed to endogenously tag *ghs2*, but only an N-terminal 3xHA fusion yielded viable colonies as the sole copy of *ghs2* at its native locus. C-terminal tagging of *ghs2* using standard protocols with GFP-*kanMX6* or mNG-*kanMX6*, with and without flexible linker sequences, failed to generate any positive integrants. N-terminal tagging by cloning *mCherry* or mNG in-frame with the *ghs2* start codon was also unsuccessful. In addition, we generated pIRT2-based constructs in which mNG was inserted internally after residues 95, 160, or 170, but none of these configurations supported integration as the sole functional copy of *ghs2*.

To generate the *3xHA-ghs2* allele we cloned gene blocks (Integrated DNA technologies) consisting of 300 bp of the 3’ UTR, 3xHA, the *ghs2^+^* CDS and *kanMX6* with 300 bp of the 5’ UTR into the BamHI site of pIRT2 using Gibson assembly. Successful cloning was confirmed by whole plasmid sequencing. Then, the full cloned sequence was amplified by PCR (5’ F oligo: TTTACGCGAAAAGTGCTTTCGAG and 3’ R oligo: ACATTCGGCTGATTGATAGATTGA) and transformed into wildtype cells using the lithium acetate method. Colonies were selected on YE agar plates containing G418 and confirmed by whole-cell PCR and sequencing from the genome.

The mCherry-Bgs3 strain was made by amplifying 300 bp of the 5’ flank, the *bgs3^+^* ORF and this fragment was combined with a gene block containing the *kanMX6* cassette and 300bp of the 3’ flank of *bgs3* using Gibson assembly into a pIRT2 vector. Next, a gene block was designed containing 300bp of the 5’ flank, the first 66 bp of *bgs3^+^*, the sequences encoding mCherry and was combined with the PCR product that included *bgs3^+^*nucleotides 67-end, *kanMX6* after the stop codon and 300 bp of the 3’ flank that were amplified from the previous pIRT2 vector. These DNA fragments we assembled into a new pIRT2 vectors with Gibson assembly. All plasmids used in this study were sequenced by Plasmidsaurus using Oxford Nanopore Technology with custom analysis and annotation. Lastly, the full *mCherry-bgs3:kanMX6* gene with 300bp flanks was amplified (5’ F oligo: CGCATTCCGGTTTCAATATACTCACA and 3’ R oligo:TGCAAGGCATGCAAATTCACAA) and transformed into wildtype cells using lithium acetate method. Colonies resistant to G418 were tested with whole-cell PCR and confirmed with microscopy. Newly generated *S. pombe* strains, plasmids, and other reagents described in this study are available from the corresponding author upon reasonable request.

For Figure 6a and 6c, a 150 mM D75-4590 (MedChemExpress, cat. # HY-134655) stock solution was made in DMSO. 1 mL of cells grown in EMM with appropriate supplements were treated with the indicated concentration of D75-4590 or DMSO vehicle control for 3h. For Figure 6d, a D75-4590 stock solution of 20 mM was made in DMSO. 1 mL of cells were spun down and resuspended in 50-100 uL of YE and then treated with DMSO control or the appropriate concentration of D75-4590. The cells were fixed with 70% ice-cold ethanol for 1 hour at 4°C, washed in PBS three times and then stained with methyl blue prior to imaging.

### Isolation of the temperature sensitive allele with error-prone PCR

The *bgs3-1* temperature sensitive allele was constructed by first tagging *bgs3^+^* with the sequences encoding *kanMX6* after the *bgs3^+^* stop codon. Next, genomic DNA from this strain was isolated and error-prone PCR was performed to introduce mutation into the *bgs3* gene. The mutations were introduced by using oligonucleotides 300 bp upstream or downstream of the *bgs3:kanMX6* ORF (5’ F oligo: CGCATTCCGGTTTCAATATACTCACA and 3’ R oligo:TGCAAGGCATGCAAATTCACAA), the EX taq polymerase (Takara, cat. # 4025), 3 mM MgCl_2_ and unbalanced dNTPs (Takara, cat. # RR01BM) where one of the nucleotides is a 10x higher concentration (100 mM versus 10 mM) than the others^94^. Subsequently, the PCR product was transformed into wildtype cells using the lithium acetate protocol^92^, grown at 25°C on non-selective YE plates for 2 days, then replica plated to YE+G418 (Geneticin, 100 µg/mL, Thermo Fisher Scientific; cat. # 11811031) and allowed to grow for one week for colonies form. Then, the plates were replica plates to YE plates containing Phloxine B (7.5 mg/mL, Millipore-Sigma, cat # P4030) at 36°C. Phloxine B stains dead cells as a dark pink color. Colonies were visually screened for dark pink colonies on the YE+Phloxine B plates and source colonies from the YE+G418 plate was obtained for further analysis. The *bgs3-1* allele was backcrossed to wildtype twice and verified by rescue of a vector carrying *bgs3^+^*. The allele was sequenced by Plasmidsaurus using Oxford Nanopore Technology with custom analysis and annotation.

### Immunoprecipitations and immunoblotting

For immunoprecipitation of GFP-Ghs2 in Fig. S1d, whole-cell lysates were prepared in NP40 buffer [6 mM Na_2_HPO_4_, 4 mM NaH_2_PO_4_, 1% NP-40, 150 mM NaCl, 50 mM NaF, 0.1 mM Na_3_VO_4_, 2 mM EDTA, 1 mM Phenylmethylsulfonyl fluoride (PMSF), 2 mM benzamidine, 0.5 mM diisopropyl fluorophosphate (DIFP), 5 μg/mL leupeptin] and protease inhibitor tablet (cOmplete Protease Inhibitor Cocktail, Roche), in native conditions as previously described^95^. 30 ODs of cells were lysed by bead disruption using a FastPrep-24 cell homogenizer (MP Biomedicals; catalogue# 116004500) for 20s. Proteins were immunoprecipitated from protein lysates using GFP-TRAP agarose beads (Proteintech; cat. # gta).

For immunoprecipitation of GFP-Bgs3, HA_3_-Ghs2 and GFP-Ghs2 and for co-immunopreciptiation experiments, a modified membrane extraction protocol was performed based on published protocols^96,97^. Cells were grown to mid-log phase (A_595_ = 0.3-0.5), and cultures corresponding to 120 units of A_595_ (approximately 10 cells) were harvested. The cells were lysed by bead disruption using a FastPrep-24 cell homogenizer for 20s at a motor speed of 4.0 m/s. Lysates were extracted with lysis buffer (50 mM Tris-HCl pH 7.4, 200 mM NaCl, 1 mM EDTA) containing protease inhibitors (cOmplete Protease Inhibitor Cocktail, 3.5 μM Benzamidine-HCl, 100 μM PMSF and 0.5 mM DIFP). After a short clearing spin of 4,500 g for 1 minute, crude membrane and soluble fractions were separated by centrifugation for 1 h at 16,000 g at 4°C. The membrane pellets were then resuspended in the lysis buffer containing 1% NP-40. Insoluble material was cleared by centrifugation at 13,000 g for 20 min at 4°C. Proteins were immunoprecipitated with anti-HA (12CA5) and protein G agarose beads (GE Healthcare; cat. # 17-1018-02) or GFP-TRAP magnetic agarose (proteintech; cat. # gtma) and washed three times with lysis buffer with NP-40 before proteins were eluted with SDS sample buffer.

For immunoblotting, proteins were resolved by 12% PAGE or 3-8% NuPAGE, transferred by electroblotting to a polyvinylidene difluoride membrane (Immobilon P; Millipore) and incubated with the set of primary antibodies indicated at 1 µg/ml. GFP was detected with a polyclonal antibody (proteintech; cat. # 50430-2-AP, RRID: AB_11042881) at a 1:1,000 dilution, RFP was detected with a monoclonal antibody (proteintech; cat. # 6g6, RRID: AB_2631395) at a 1:2,000 dilution, HA was detected with a monoclonal antibody 12CA5 (Vanderbilt Antibody and Protein Resource; cat. # Anti-HA 12CA5, RRID: AB_2923038) at a 1:1,000 dilution and - tubulin was detected with a monoclonal antibody (Sigma-Aldrich; cat. # T6557, RRID: AB_477584) at a 1:10,000 dilution. Primary antibodies were detected with secondary antibodies coupled to Alexa Fluor 680 goat anti-mouse (LI-COR Biosciences; cat. # 926-68020, RRID: AB_10706161) or goat anti rabbit (LI-COR Biosciences; cat. # 926-68021, RRID: AB_10706309) or IRDye800 goat anti-mouse (LI-COR Biosciences; cat. # 926-32210, RRID: AB_621842) or goat anti-rabbit (LI-COR Biosciences; cat. # 926-32211, RRID: AB_621843) all at a 1:10,000 dilution and visualized using an Odyssey Infrared Imaging System (LI-COR Biosciences).

### Microscopy

Live-cell and fixed-cell images of *S. pombe* cells were acquired using a Zeiss Axio Observer inverted epifluorescence microscope with Zeiss 63× oil (1.46 NA) and captured using Zeiss ZEN (V3.0, Blue edition) software and Axiocam 503 monochrome camera (Zeiss).

Intensity measurements were made with ImageJ software (V2.16.0) (http://rsweb.nih.gov/ij/)^98^. For all intensity measurements, the background was subtracted by creating an ROI in the same image where there were no cells (Waters, 2009). Measurements of protein localization were done in 3 biological replicates and images were acquired using Z-series optical sections of 0.5 μm spacing for a total of 4.5-5.0 μm. Images for quantification were not deconvolved and sum projected. Cells were fixed with 70% ethanol for DAPI and methyl blue (MB) staining as described previously^99^.

### Transmission electron microscopy

*S. pombe* strains were grown to mid-log phase (A_595_ ≈ 0.3) and were harvested by centrifugation at 1000 × g for 5 minutes. Samples were loaded into 3 mm high-pressure freezer specimen carriers (Ted Pella) and vitrified using a Leica EM ICE high-pressure freezer. Samples were freeze-substituted in 1.5% uranyl acetate in methanol at −90°C for 48 hours followed by infiltration HM20 Lowicryl for 48 hours at −45°C. The resin was UV polymerized at −45°C for 24 hours, followed by an additional 24 hours of UV at 0°C. The samples were cut on a Leica Enuity at a nominal thickness of 70 nm and collected on either 300 mesh Ni grids or Cu slot grids with a carbon film. Grids were stained with 1% phosphotungstic acid, 2% uranyl acetate, and lead citrate to contrast the cell walls. Samples were imaged by transmission electron microscopy (TEM) on a JEOL 2100+ TEM operating at 200 keV using an AMT nanosprint 15mkII CMOS camera. Cell wall thickness was measured using ImageJ (V1.8.0).

Measurements were obtained from 10 individual cells, with 10 cell wall thickness (non-deposit) and 8 individual cells, with 5 cell wall thickness (deposit) readings recorded per cell. Differences in thickness among wildtype, *bgs3-1*, and *ghs2-2* were evaluated using one-way single factor ANOVA (\*\*\*\**p* < 0.0001) and *t*-test with one tail (\*\**p* ≤ 0.01). Data analysis was carried out in Microsoft Excel 365 (V16.111.3), and violin plots were created using OriginPro (V9.0).

### AlphaFold3 structural prediction and sequence alignment

Protein structure predictions were generated with the AlphaFold3 server (https://alphafoldserver.com)^38^. Protein sequences from Pombase were input into the server with a single copy of each protein^100^. Full-length Bgs3 (Q9P377) and Ghs2 (O13789) or full-length Bgs1 (Q10287) and Sbg1 (O13619) were modeled together. The automatic seed setting was used for all predictions and the top-ranked “model_0” prediction was selected to extract interface information and visualization using the PyMOL molecular graphics system (V3.1.4.1, Schrodinger, LLC). The Ghs2-Bgs3 model was generated on July 21^st^, 2025 using seed 361330275; The Sbg1-Bgs1 model was generated on May 11^th^, 2024 using seed 815971004. Protein sequence alignment was performed using Clustal Omega^80,101^.

### Solid-state NMR spectroscopy

Cells were grown to mid-log phase (A_595_ = 0.3-0.5) and were harvested by centrifugation at 1000 × g for 5 minutes. The resulting cell pellets were washed thoroughly with Milli-Q water to remove small water-soluble molecules and excess ions, ensuring retention of natural cellular hydration. Approximately 20 mg (wildtype, *bgs3-1, ghs2-2*) and 10 mg (*bgs3-1 ghs2-2*) of the prepared cell material was packed into 3.2 mm magic-angle spinning (MAS) rotors (HZ16916, Cortecnet) for subsequent solid-state NMR analysis.

The EMM media used for labeling *S. pombe* contains ^13^C-labeled D-glucose (10.0 g/L, CLM-1396-PK, Cambridge Isotope Laboratories) 5g/L ammonium chloride, 3.0 g/L potassium hydrogen phthalate, 2.2 g/L sodium hydrogen phosphate, 20.0 mL/L salt solution (52.5 g/L magnesium dichloride hexahydrate, 0.735 g/L calcium dichloride dihydrate, 50.0 g/L potassium chloride, and 2.0 g/L sodium sulfate), 1.0 mL/L vitamin solution (1.0 g/L pantothenic acid, 10.0 g/L nicotinic acid, 10.0 g/L inositol, 10.0 mg/L biotin), 0.1 mL/L mineral solution (5.0 g/L boric acid, 4.0 g/L manganese sulfate, 4.0 g/L zinc sulfate heptahydrate, 2.0 g/L ferrous chloride hexahydrate, 0.4 g/L molybdic acid, 1.0 g/L potassium iodide, 10.0 g/L citric acid, and 0.4 g/L copper sulfate pentahydrate), along with 225 mg/L adenine, 225 mg/L uracil, and 225 mg/L leucine. The solution was autoclaved prior to use.

NMR experiments were conducted on Bruker Avance Neo spectrometer at MSU Max T. Roger facility, with ^1^H Larmor frequency of 800 MHz (18.8 Tesla). All experiments were collected using 3.2-mm MAS HCN probes under 15 kHz MAS at 293 K. ^13^C chemical shifts were externally referenced to the adamantane CH_2_ peak at 38.48 ppm on the tetramethyl silane (TMS) scale. Typical radiofrequency field strengths, unless specifically mentioned, were 80-100 kHz for ^1^H decoupling, 62.5 kHz for ^1^H hard pulses, and 50-62.5 kHz for ^13^C. The experimental parameters are summarized in Supplementary Table 2.

Solid state NMR experiments were carried out using 1D and 2D experimental methods.

1D ^13^C spectra were measured using different polarization methods to selectively detect the rigid and mobile components. Rigid components were characterized by employing a dipolar based ^1^H-^13^C cross-polarization (CP) with a 1-ms contact time, adhering to Hartmann-Hahn match conditions of 62.5 kHz for ^1^H and 47.5 kHz for ^13^C channels (Supplementary Fig. 5). Mobile components were measured by 1D ^13^C direct polarization (DP) using short (2 s) recycle delays (Supplementary Fig. 6). Extending the recycle delays to 35 s allowed us to achieve quantitative detection of all molecules through the same ^13^C DP experiment (Supplementary Fig. 7). Resonance assignment of rigid molecules was achieved through the measurement of 2D ^13^C-^13^C 53-ms CORD homonuclear correlation spectra^102^, which detect intramolecular cross-peaks. For mobile molecules, 2D DP refocused J-INADEQUATE spectra were measured to track through bond connectivity, with each of the four tau delays optimized to 2.3-ms^103^. The resolved chemical shifts were compared with the values indexed in the Complex Carbohydrate Magnetic Resonance Database (CCMRD) to validate the chemical nature of the carbohydrates^104^. The ^13^C chemical shifts of carbohydrates identified in *S. pombe* have been tabulated in Supplementary Table 3. using Topspin 4.3.0 and all the graphs were generated through OriginPro 9.0. Illustrative Figs were prepared using Adobe Illustrator 2023 V27.2.0. The molecular composition was estimated by analyzing the intensities of distinct carbohydrate signals observed in 2D ^13^C spectra^105^. Rigid molecules were analyzed using CORD spectra^102^, while mobile molecules were evaluated using DP refocused J-INADEQUATE spectra. The estimation was based on peak volumes, with only well-resolved signals included in the analysis. The relative abundance of each polysaccharide was quantified by normalizing the summation of integrated peak volumes by their respective cross-peak counts. The standard error for each polysaccharide was calculated by dividing the standard deviation of the integrated peak volumes by the total cross-peak count. The overall standard error was determined as the square root of the sum of the squared standard errors for all polysaccharides. To estimate the percentage error for each polysaccharide, the standard error of the specific polysaccharide was divided by its average integrated peak volume. This value was then multiplied by the ratio of the polysaccharide’s standard error to the total integrated peak volume, adjusted by its relative abundance (Supplementary Table 4, 5).

### Susceptibility Assays

Susceptibility assays were performed based on previous studies^106^. D75-4590 (MedChemExpress, cat. # HY-134655) was kept at a 150 mM stock solution in DMSO and used at the indicated final concentrations in the relevant tables and figures. For determining the minimum inhibitory concentration (MIC)-2 (the lowest drug concentration showing 50% growth inhibition compared to the control without drug^74^), log-phase cultures grown in EMM supplemented with 225 mg/L adenine, 225 mg/L uracil, and 225 mg/L leucine and were diluted to a cell density of 1-3 x 10^6^ cells/mL in media containing increasing concentrations of D75-4590 (0.125-4 mM) or an equal volume of solvent (0.65-2.6% DMSO), which was the control cell culture. The 400 μL cell cultures were grown in 48 well plates on an orbital roller at 29°C. Turbidity was examined after 24 h of growth with a Biotek Synergy HT microplate reader at A_595_. The absorbance data was represented as: −(0-4.9% turbidity), + (5-9.9% turbidity), ++ (10-24.9% turbidity), +++ (25-49.9% turbidity), ++++ (50-74.9% turbidity), and +++++ (75-100% turbidity). The values calculated and reported in Table 1 are from at three independent experiments.

### Statistical analysis

All statistical analyses were performed in Prism 11.0.2 (Graphpad software). No data were excluded from the analysis.

## Data availability

The data generated in this study underlying Figures 1-4 and 6-7 and Supplementary Figures 1-4 and 9-10 as well as Table 1 have been deposited in the Mendeley Data database under accession codes <u>10.17632/8j46dhsyxr.1</u> and <u>10.17632/y5v9w2jd6k.1</u>. The solid state NMR data underlying Figure 5 and Supplementary Figures 5-8 have been deposited in Zenodo and under accession code <u>10.5281/zenodo.20836992</u>. Source Data are provided as a source data file.

## Acknowledgements

We thank Dr. Evan Krystofiak and Rachel Hart for their help with the EM imaging.

## Funding

K.L.G. discloses support for the research of this work from National Institutes of Health (NIH) [grant number R35GM131799]. T.W. discloses support for the research of this work from NIH [grant number R01AI173270]. The Vanderbilt Cell Imaging Shared Resource core discloses support for the research of this work from NIH [grant number CA68485], [grant number R24OD037694], [grant number S10MH137068] and [grant number S10OD034315].

## Author contributions

A.H. Willet and K.L. Gould conceived the study. A.H. Willet performed biochemical experiments, AlphaFold3 modelling and imaging experiments. L.A. Turner performed the cell growth assays and generated the *bgs3-1* allele. A. Jacob and A.K.A. Alsanad conducted solid-state NMR experiments and data analysis. A.H. Willet and K.L. Gould wrote the original draft of the manuscript and all authors edited and revised the final manuscript. K.L.G. and T.W. supervised the project.

## Competing interests

The authors declare no competing interests.

## Notes

### Competing Interest Statement

The authors have declared no competing interest.

### Summary of Updates

Figure 2 revised, Figure 5 revised, Figure 6 revised, Figure 7 revised, Supplemental files updated.

