## Supplementary data for "A Bipartite Glucan Synthase-Remodeler Module Organizes Branched Glucan Assembly in the Fungal Cell Wall"

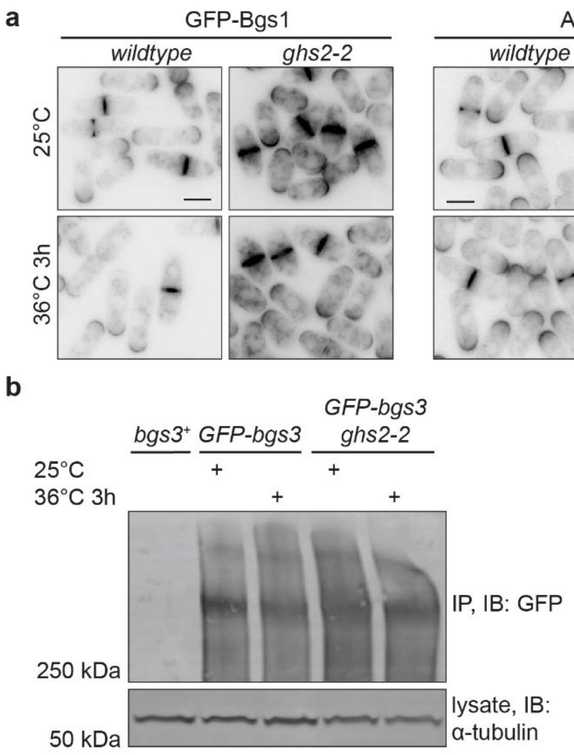


**Supplementary Figure 1. Bgs1, Bgs4 and Ags1 localization is unperturbed in *ghs2-2* cells.** a) Representative live-cell images of GFP-Bgs1, Ags1-RFP or GFP-Bgs4 in wildtype or *ghs2-2* cells. The strains were grown at 25°C and then shifted to 36°C for 3 hours. Images were acquired at both temperatures. b) GFP-Bgs3 was immunoprecipitated from the indicated cell lysates with GFP-Trap beads and incubated at 4°C for 1 h. Bound protein was resolved by SDS-PAGE and visualized by Western blot. The blot shown is representative of 2 independent replicates. α-tubulin was used as a lysate control (bottom). Scale bars, 5 µm.

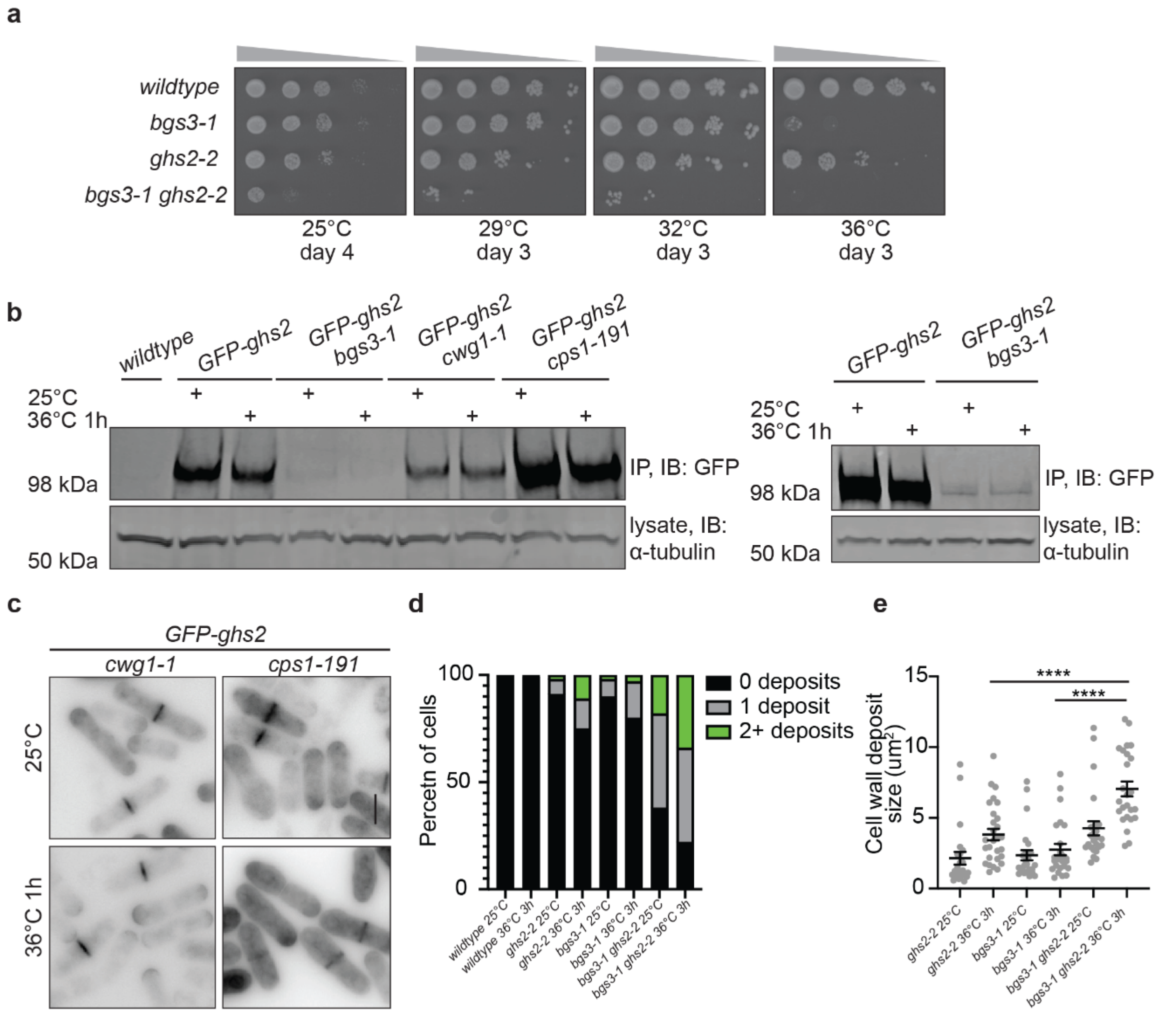


**Supplementary Figure 2. *bgs3-1* and *ghs2-2* are synthetically sick.** a) 10-fold serial dilutions of the indicated strains grown at the indicated temperatures. b) GFP-Ghs2 was immunoprecipitated from the indicated cell lysates with GFP-Trap beads and incubated at 4°C for 1 h. Bound protein was resolved by SDS-PAGE and visualized by Western blot. The blots shown is representative of 2 independent replicates. The right blot has double the amount of protein loaded than the left blot to better visualize the GFP-Ghs2 levels in *bgs3-1* cells. α-tubulin was used as a lysate control (bottom). c) Representative images of *cps1-191* or *cwg1-1* cells expressing GFP-Ghs2 from the inducible *nmt81* promoter for 24 hours prior to imaging at 25°C. The cells were then shifted to 36°C for 1 hour and imaged again. Scale bar, 5 µm. d) Quantification of the percent of cells with no abnormal cell wall deposits, 1 abnormal cell wall deposit of 2 or more abnormal cell wall deposits. n = 309 for *ghs2-2* 25°C, n = 300 for *ghs2-2* 36°C, n = 319 for *bgs3-1* 25°C, n = 349 for *bgs3-1* 36°C, n = 304 for *ghs2-2 bgs3-1* 25°C and n = 320 for *ghs2-2 bgs3-1* 36°C. e) Quantification of the size of the cell wall deposits in the indicated strains and conditions.

n = 23 for *ghs2-2* 25°C, n = 28 for *ghs2-2* 36°C and n=25 for both *bgs3-1* and *ghs2-2 bgs3-1* conditions. Error bars represent mean ± SEM. Statistical analysis was performed using Welch's *t*-test. ****$P$<0.0001.

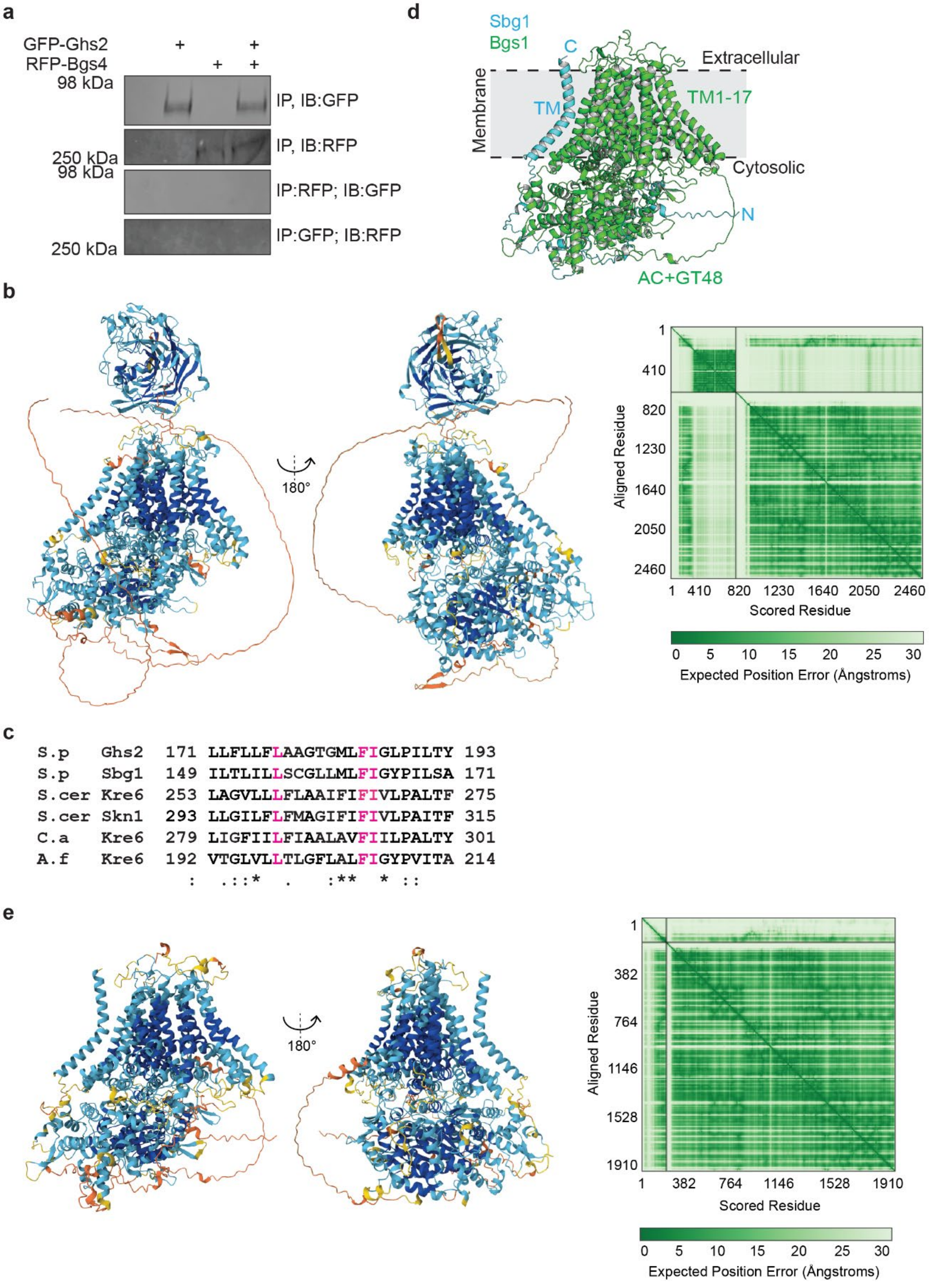


**Supplementary Figure 3. The Sbg1-Bgs1 complex has a similar architecture to the Ghs2-Bgs3 complex.** a) Anti-GFP and anti-RFP immunoprecipitations from the indicated strains. Both immunoprecipitations were blotted for with anti-GFP and anti-RFP antibodies. b) Left, the AlphaFold3 predicted model of the Ghs2-Bgs3 complex from Figure 3c colored by the predicted local difference distance test (pLDDT). Right, the predicted aligned error plot for the AlphaFold3 generated structural prediction. c) Sequence alignment of the transmembrane helix sequences from *S. pombe* (S.p) Ghs2 and Sbg1, *S. cerevisiae* (S.cer) Kre6 and Skn1, *Candida albicans* (C.a) Kre6 and *Aspergillus fumigatus* (A.f) Kre6. The residues in magenta are the same in all proteins and mutated in the Ghs2-3A protein depicted in f. "*" denotes the same residue, ":" denotes strongly similar residues and "." denotes weakly similar residues. d) AlphaFold3-generated model of Bgs1-Sbg1 complex. Bgs1 is in green and Sbg1 is in cyan. e) Left, the AlphaFold3 predicted model of the Sbg1-Bgs1 complex from d, colored by the predicted local difference distance test (pLDDT). Right, the predicted aligned error plot for the AlphaFold3 generated structural prediction. b, e) Dark blue pLDDT indicates very high confidence and a score greater than 90, cyan indicates a confident score of 90-70, yellow indicates a low score of 70-50 and orange indicates a very low score that is less than 50[1].

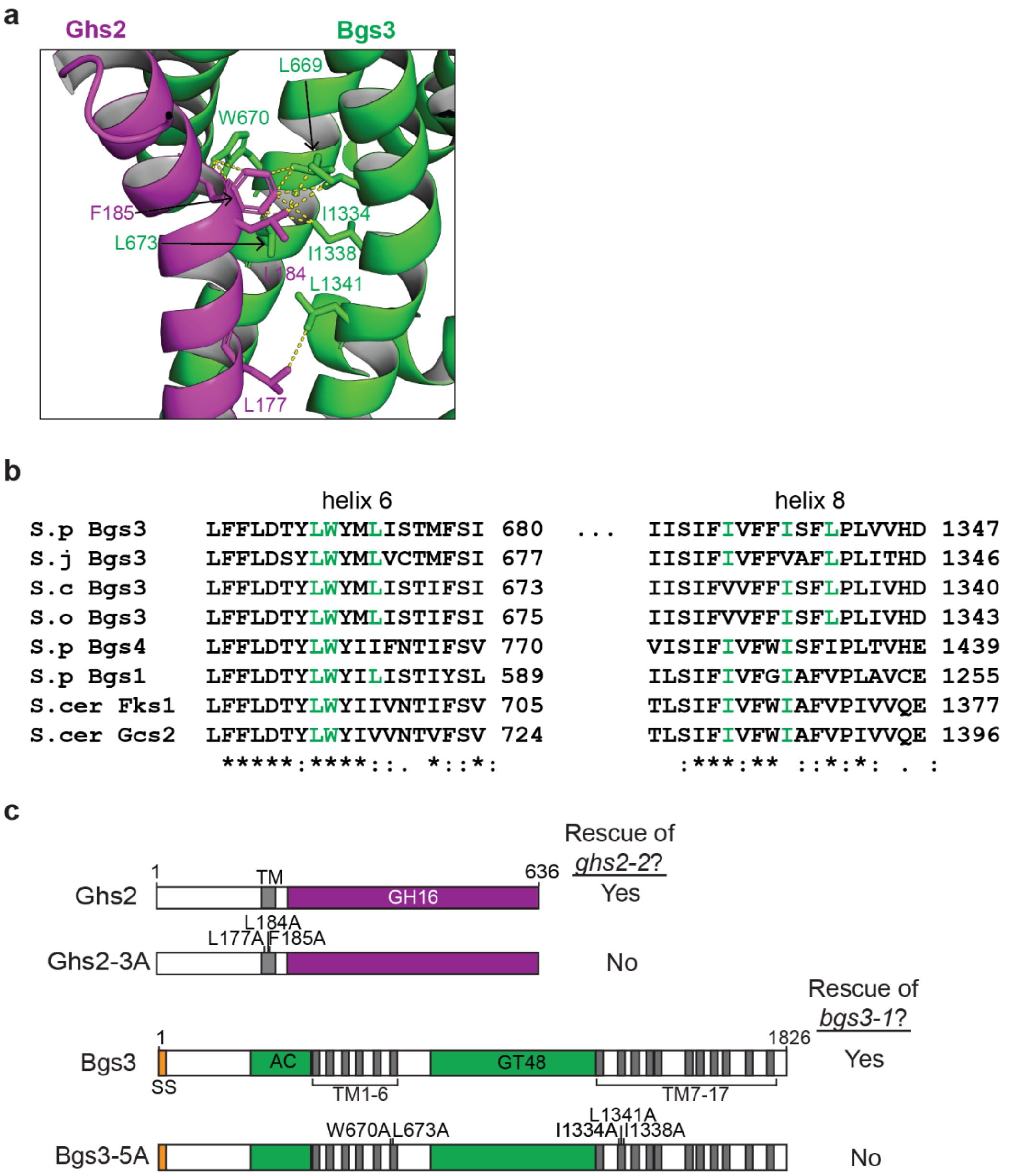


**Supplementary Figure 4. The Ghs2 and Bgs3 residues predicted to interact within the transmembrane helices.** a) Zoomed-in view of one of the AF3 predicted Ghs2-Bgs3 interaction interfaces from Figure 3c. This interaction interface is mediated by the Ghs2 transmembrane helix and the Bgs3 transmembrane helices 6 and 8. Ghs2 is in magenta and Bgs3 is in green. The amino acids predicted to mediate the interaction are shown as sticks with side chains and are labeled. b) Sequence alignment of the indicated amino acid sequences of the predicted helix 6 and 8 of the indicated β-glucan synthases. *S. pombe* (S.p), *S. cerevisiae* (S.cer), *Schizosaccharomyces japonicus* (S.j),

*Schizosaccharomyces cryophilus* (S.c) and *Schizosaccharomyces octosporus* (S.o). The residues in green are mutated in the Bgs3-5A protein depicted in f. "*" denotes the same residue, ":" denotes strongly similar residues and "." denotes weakly similar residues. c) Schematics, drawn to scale, of the indicated proteins. Transmembrane (TM) helices are in gray, the accessory domain (AC) and glycosyl transferase 48 (GT48) domains are in green, and the glycoside hydrolase 16 (GH16) domain is in purple and the signal sequence (SS) is in orange. *ghs2* was expressed from the *nmt1* promoter and *bgs3* was expressed from the endogenous promoter. The plasmids containing the genes encoding the indicated proteins were transformed into *ghs2-2* or *bgs3-1* cells, grown at 25°C and then struck out at 36°C and incubated for 24-48h to test for rescue.

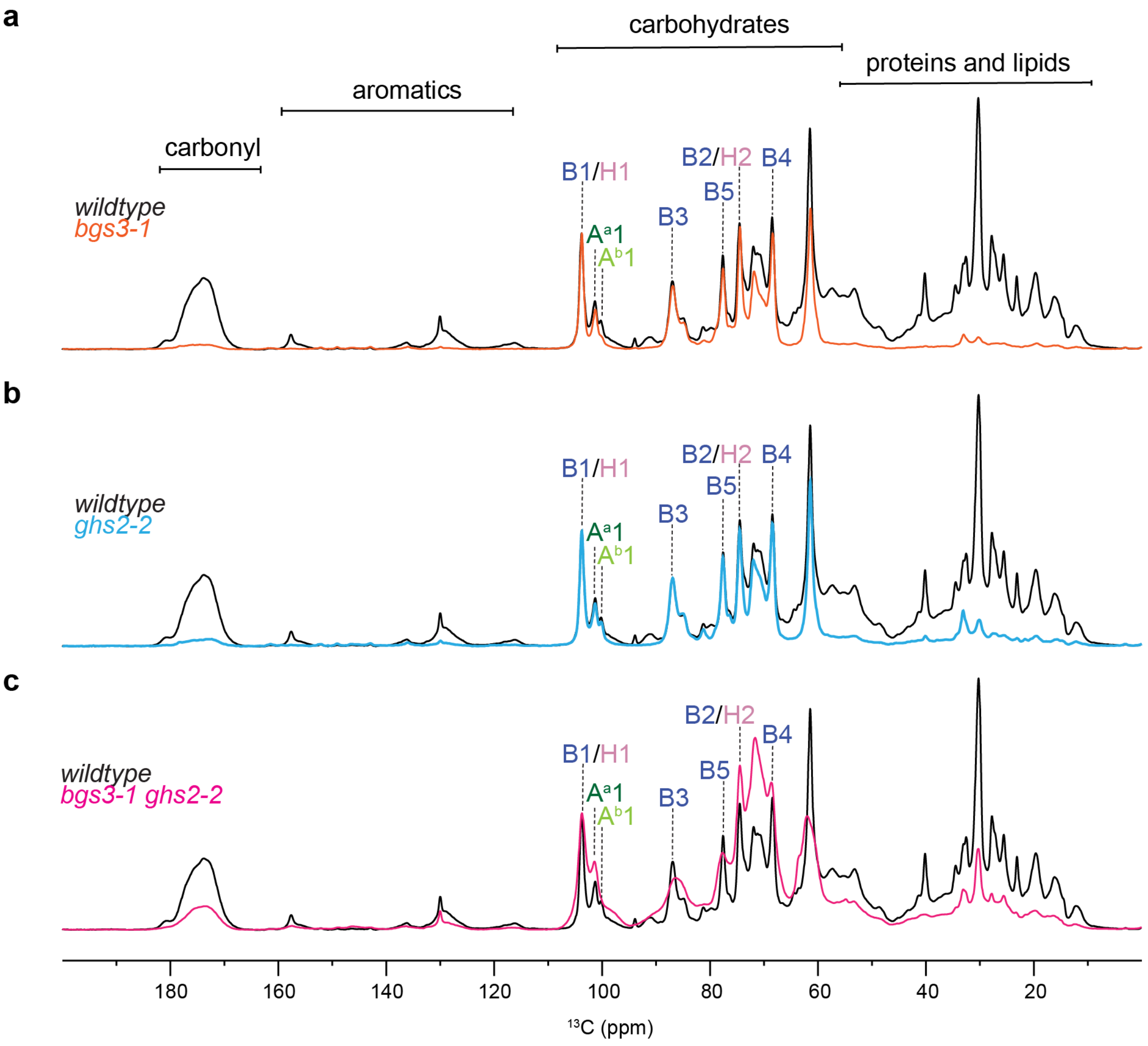


**Supplementary Figure 5. Comparison of rigid polysaccharide components in the cell wall.** 1D $^{13}C$ CP spectra comparing the rigid polysaccharide composition of the wildtype strain (black) with a) *bgs3-1* (orange), b) *ghs2-2* (cyan) and c) *bgs3-1 ghs2-2* (pink). Abbreviations: B, β-1,3-glucan; H, β-1,6-glucan; A, α-1,3-glucan.

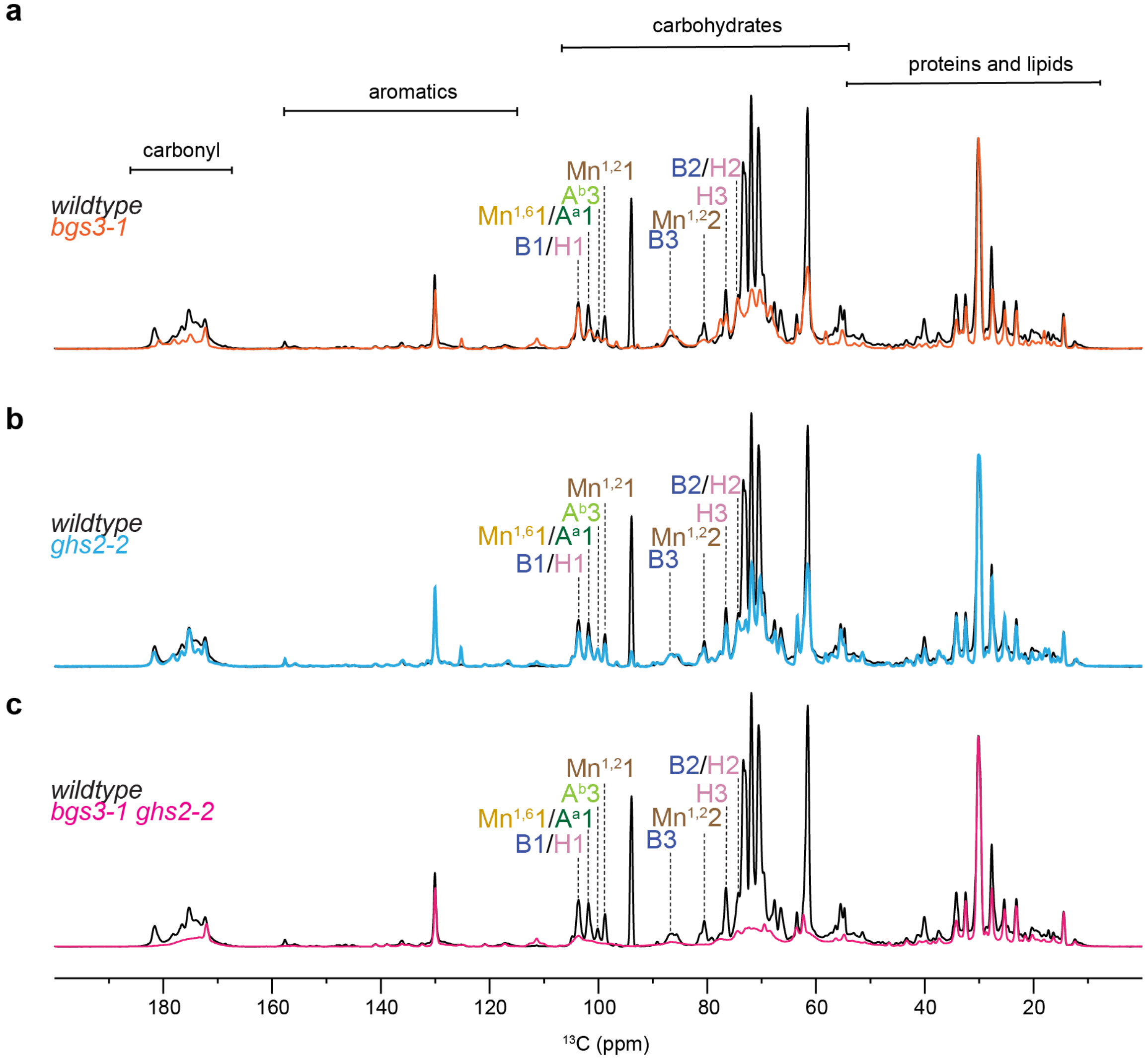


**Supplementary Figure 6. Comparison of mobile polysaccharide components in the cell wall.** 1D $^{13}C$ DP spectra (with a short recycle of 2 s) spectra comparing the mobile polysaccharide composition of the wildtype strain (black) with a) *bgs3-1* (orange), b) *ghs2-2* (cyan) and c) *bgs3-1 ghs2-2* (pink). Abbreviations: B, β-1,3-glucan; H, β-1,6-glucan; A, α-1,3-glucan; $Mn^{1,2}$, α-1-2-mannose; $Mn^{1,6}$, α-1,6-mannan.

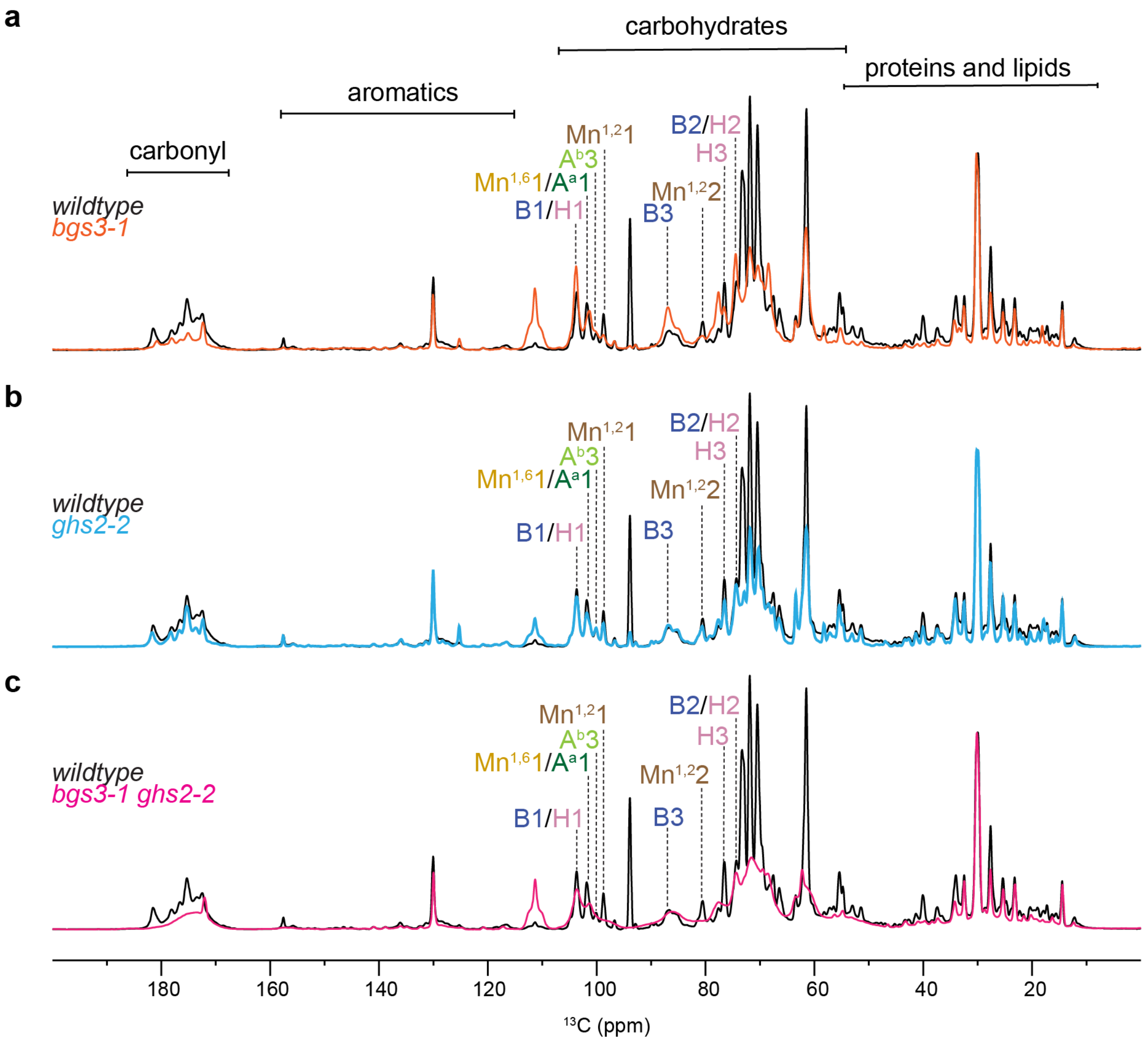


**Supplementary Figure 7. Quantitative analysis of polysaccharide components in the cell wall.** 1D $^{13}$C DP spectra (with a long recycle delay of 35 s) enabling quantitative detection of all polysaccharides. Spectra compare the polysaccharide composition of the wildtype strain (black) with a) *bgs3-1* (orange), b) *ghs2-2* (cyan) and c) *bgs3-1 ghs2-2* (pink). Abbreviations: B, β-1,3-glucan; H, β-1,6-glucan; A, α-1,3-glucan; $Mn^{1,2}$, α-1-2-mannose; $Mn^{1,6}$, α-1,6-mannan.

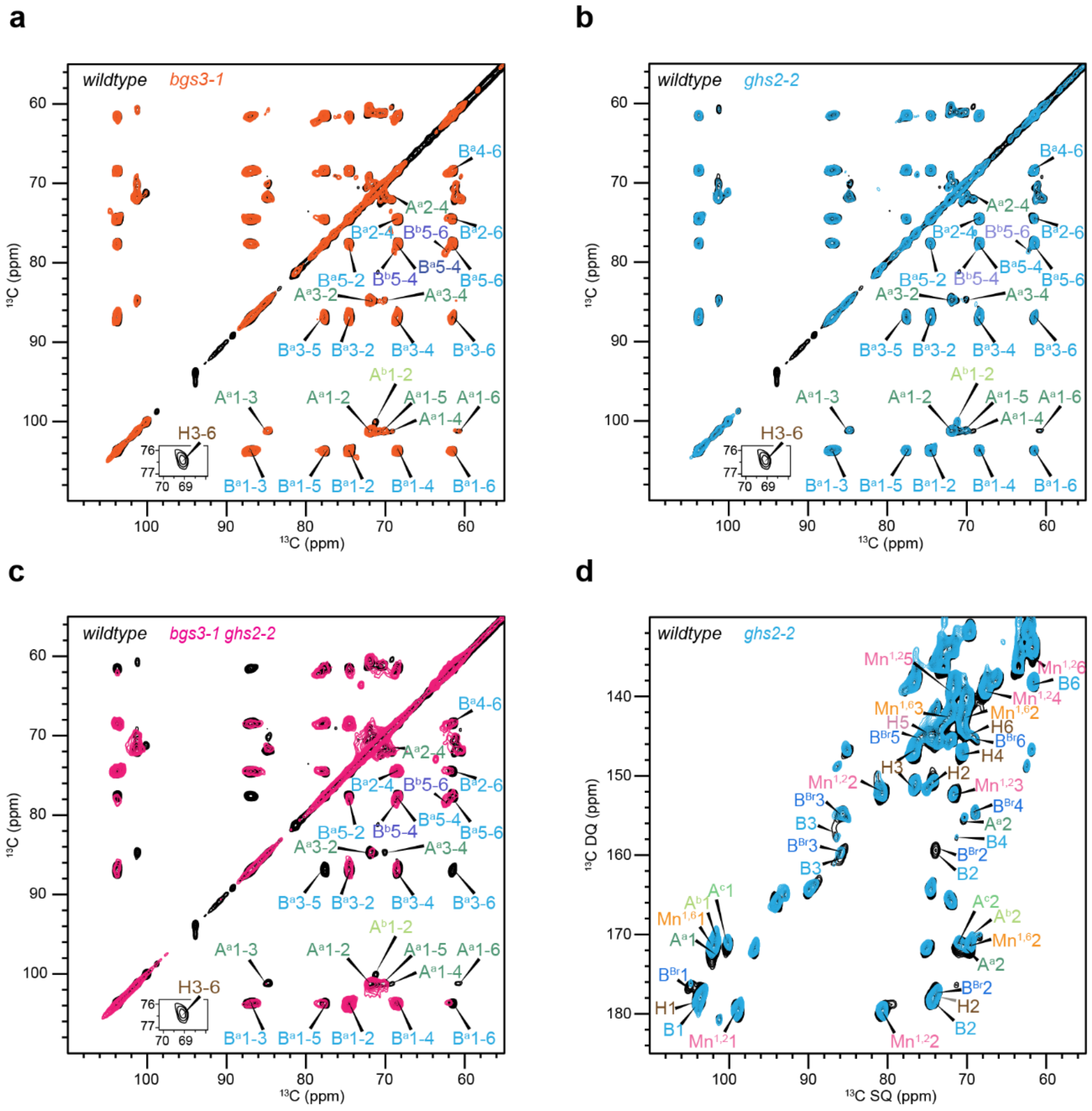


**Supplementary Figure 8. 2D ssNMR analysis of polysaccharide components in the cell wall.** 2D $^{13}$C-$^{13}$C 53-ms CORD spectrum resolving all carbons sites of rigid polysaccharides in the cell walls of wildtype (black) compared with a) *bgs3-1* (orange), b) *ghs2-2* (blue) c) *bgs3-1 ghs2-2* (pink). b) Carbohydrate region of 2D $^{13}$C DP refocused J-INADEQUATE spectra. Mobile molecules are detected in wildtype (black) and *ghs2-2* (blue). Abbreviations: B, β-1,3-glucan; H, β-1,6-glucan; $B^{Br}$, β-1,3,6-glucan; A, α-1,3-glucan; $Mn^{1,2}$, α-1-2-mannose; $Mn^{1,6}$, α-1,6-mannan.

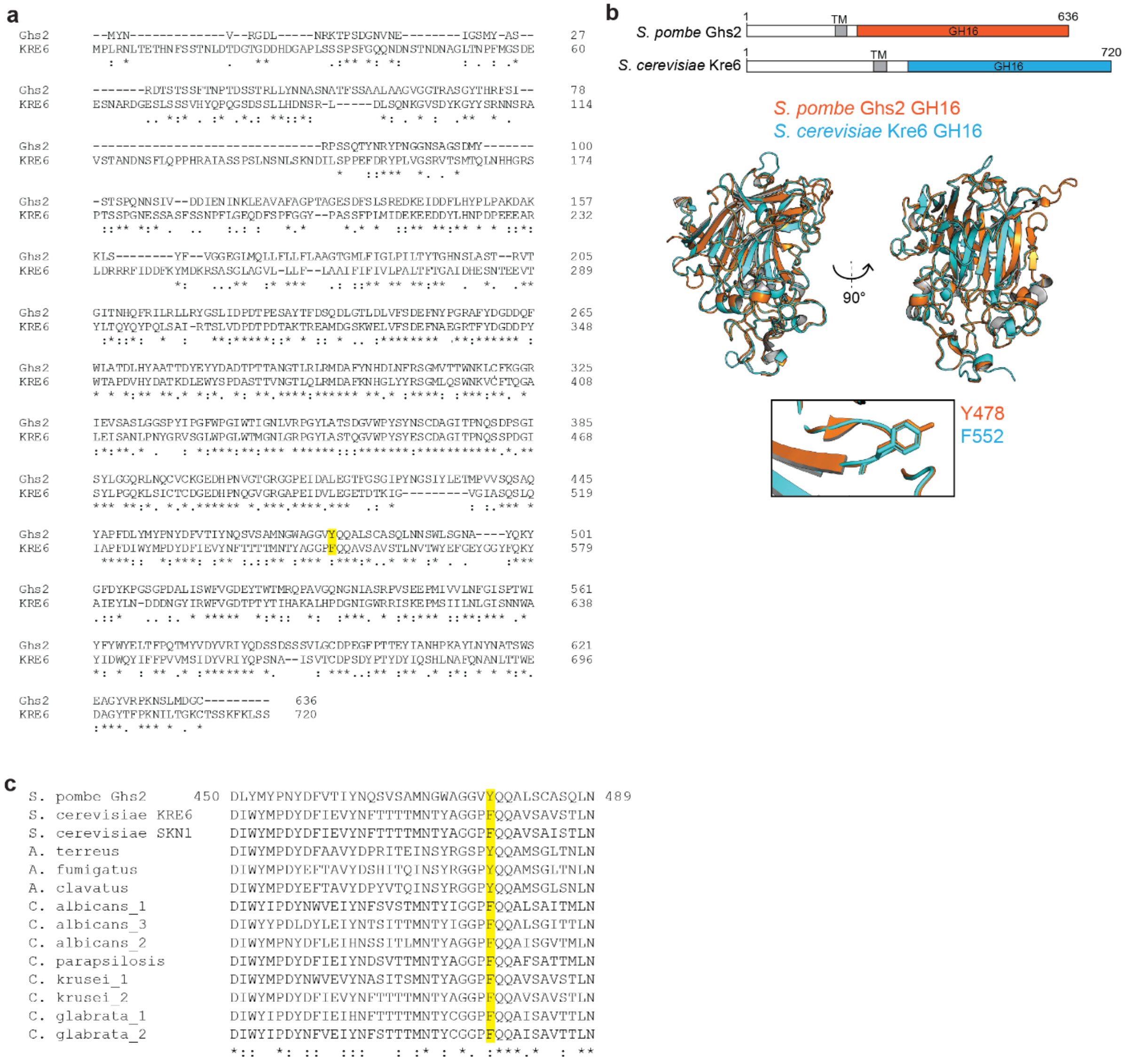


**Supplementary Figure 9. Comparison of *S. pombe* Ghs2 and *S. cerevisiae* Kre6.** a) Sequence alignment of the indicated Ghs2 amino acids and the listed homologs. F478 and the conserved corresponding F or Y residues are highlighted in yellow. b) Protein sequence alignment for full-length *S. pombe* Ghs2 and *S. cerevisiae* Kre6. For a and b "*" denotes the same residue, ":" denotes strongly similar residues and "." denotes weakly similar residues. c) Schematics, drawn to scale, of the indicated proteins and their domains. Transmembrane (TM), Glycoside hydrolase 16 (GH16). Below are the aligned AlphaFold-predicted GH16 domains of Ghs2 (orange) and Kre6 (cyan). The bottom zoomed-in view shows the predicted position of the side chains of the labelled residues.

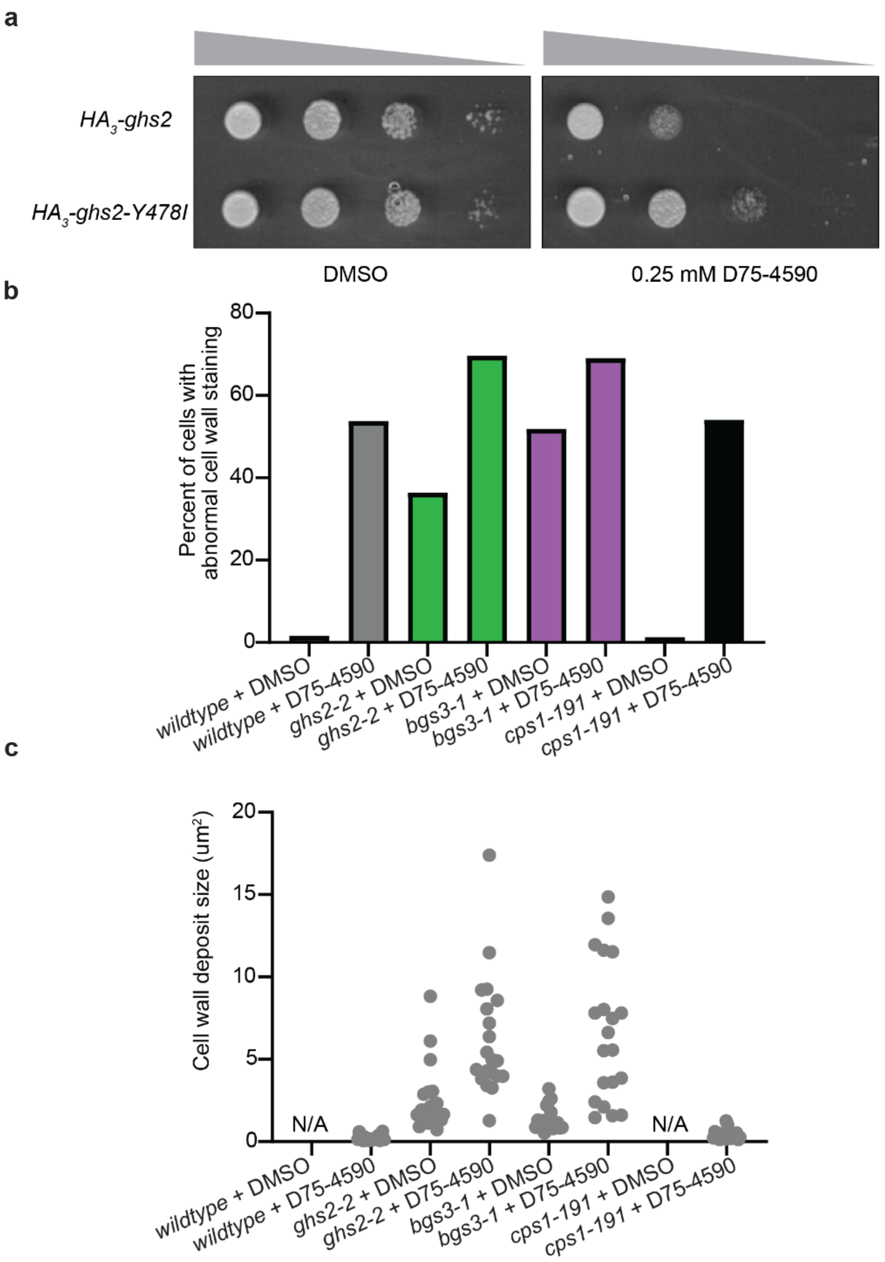


**Supplementary Figure 10. Comparison of strain phenotypes when treated with D75-4590.** a) 10-fold serial dilutions of the indicated strains grown at 29°C for 3 days. b) Quantification of the percent of cells with abnormal cell wall staining from cells in Figure 6d. Wildtype is in gray, *ghs2-2* is in green, *bgs3-1* is in purple and *cps1-191* is in black. n = 310 for WT DMSO, n = 342 for WT D75-4590, n = 318 for *ghs2-2* DMSO, n = 420 for *ghs2-2* D75-4590, n = 335 for *bgs3-1* DMSO, n = 353 for *bgs3-1* D75-4590, n = 313 for *cps1-191* DMSO and n = 316 for *cps1-191* D75-4590 from two independent experiments. c) Quantification of the size of cell wall deposits from Figure 6d. n = 20 for each from two independent experiments. No deposits were present in the wildtype and *cps1-191* DMSO treated cells.

**Supplementary Table 1. *S. pombe* strains used in this study.**

| Strain | Genotype | Source |
|---|---|---|
| **Figure 1** | | |
| KGY11110 | *bgs4Δ::ura4$^{+}$ GFP-bgs4:leu1$^{+}$ leu1-32 ura4-D18 his3-D1 h$^{-}$* | Cortès et al., 2005[2] |
| KGY2350-2 | *bgs3Δ::ura4$^{+}$ GFP-bgs3:leu1$^{+}$ leu1-32 ura4-D18 his3-D1 h$^{+}$* | Cortès et al., 2007[3] |
| KGY2098-2 | *ghs2-2:kanMX6 bgs4Δ::ura4$^{+}$ GFP-bgs4:leu1$^{+}$ leu1-32 ura4-D18 h$^{+}$* | This study |
| KGY1534-2 | *ghs2-2:kanMX6 bgs3Δ::ura4$^{+}$ GFP-bgs3:leu1$^{+}$ leu1-32 ura4-D18 h$^{+}$* | This study |
| KGY246 | *ura4-D18 leu1-32 ade6-M210 h$^{-}$* | Lab stock |
| KGY3536-2 | *bgs3-1:kanMX6 ura4-D18 leu1-32 ade6-M210 h$^{-}$* | This study |
| KGY16854 | *cwg1-1 ura4-D18 leu1-32 ade6-M210 h$^{+}$* | Ribas et al., 1991[4] |
| **Figure 2** | | |
| KGY6297-2 | *ghs2-2:kanMX6 ura4-D18 leu1-32 ade6-M210 h$^{+}$* | Lab stock |
| KGY3157-2 | *ghs2-2:kanMX6 bgs3-1:kanMX6 ura4-D18 leu1-32 ade6-M210 h$^{-}$* | This study |
| **Figure 3** | | |
| KGY3167-2 | *mCherry-bgs3:kanMX6 ura4-D18 leu1-32 ade6-M210 h$^{-}$* | This study |
| KGY3973-2 | *3xHA-ghs2:kanMX6 ade6-M210 leu1-32 ura4-D18 h$^{-}$* | This study |
| KGY2520-2 | *3xHA-ghs2:kanMX6 bgs3Δ::ura4$^{+}$ GFP-bgs3:leu1$^{+}$ leu1-32 ura4-D18 ade6-M210 h$^{+}$* | This study |
| **Figure 4** | | |
| KGY3723-2 | *ghs2-2:kanMX6 mCherry-bgs3:kanMX6 ura4-D18 leu1-32 ade6-M210 h$^{-}$* | This study |
| **Figure 7** | | |
| KGY2975-2 | *3xHA-ghs2-Y478I:kanMX6 ade6-M210 leu1-32 ura4-D18 h$^{-}$* | This study |
| **Figure S1** | | |
| KGY3673-2 | *GFP-bgs1:leu1$^{+}$ bgs1Δ::ura4$^{+}$ ura4-D18 leu1-32 his3-D1 h$^{-}$* | Cortès et al., 2002[5] |
| KGY1937-2 | *ags1Δ::ura4$^{+}$ ags1-RFP:leu1$^{+}$ ade6-M210 h$^{+}$* | Cortès et al., 2012[6] |
| KGY2105-2 | *ghs2-2:kanMX6 GFP-bgs1:leu1$^{+}$ ura4-D18 leu1-32 ade6-M210 h$^{-}$* | This study |
| KGY1965-2 | *ghs2-2:kanMX6 ags1Δ::ura4$^{+}$ ags1-RFP:leu1$^{+}$ ade6-M210 h$^{+}$* | This study |
| KGY2153 | *cps1-191 ura4-D18 ade6-M210 lys1-131 h$^{-}$* | Liu et al., 1999[7] |
| **Figure S3** | | |
| KGY7559-2 | *RFP-bgs4:leu1$^{+}$ bgs4::ura4$^{+}$ leu1-32 ura4-D18 his3-D1 h$^{-}$* | Cortès et al., 2005[2] |

**Supplementary Table 2. Solid-state NMR experiments and parameters.** To be quantitative, direct pulse (DP) experiments with 35 s long recycling delay were used. cross-polarization (CP), most rigid molecules. With DP and a shorter recycling delay of 2 seconds, suppress the rigid molecules from the spectra, and with Insensitive Nuclei Enhanced by Polarization Transfer (INEPT) the most mobile molecules were selected. For 2D $^{13}C$-$^{13}C$ correlation experiments allowed to resolve rigid intramolecular peaks. 2D DQ-SQ DP J-INADEQAUTE spectra was used to detect through-bond correlations. The experimental parameters include the $^{1}H$ Larmor frequency, total experiment time (t), recycle delay (d1), number of scans (NS), The number of points for the direct (td2) and indirect (td1) dimensions, the acquisition time of the direct dimension (aq2) and the evolution time of indirect dimension (aq1), spectral width (sw1 and sw2), mixing time ($t_m$), increment delay (IN_F). The processing parameters include the window function and associated parameters.

| | **Acquisition parameters** | | | | | | | | | | | | **Processing parameters** | |
|---|---|---|---|---|---|---|---|---|---|---|---|---|---|---|
| **Experiment** | $\omega_{0,\,1H}$ (MHz) | t (h) | d1 (s) | NS | td2 | td1 | aq2 (ms) | aq1 (ms) | sw2 (ppm) | sw1 (ppm) | $t_m$ (ms) | IN_F (µs) | Window function | Parameter |
| 1D $^{13}C$ CP | 800 | 0.5 | 2.0 | 1024 | 3200 | | 16.0 | | 496.8 | | 1.0 | | GM | LB-10, GB0.05 |
| 1D $^{13}C$ DP | 800 | 0.2 | 2.0 | 512 | 3200 | | 16.0 | | 496.8 | | | | GM | LB-10, GB0.05 |
| 1D $^{13}C$ DP | 800 | 2.5 | 35.0 | 512 | 3200 | | 16.0 | | 496.8 | | | | GM | LB-10, GB0.05 |
| 2D $^{13}C$-$^{13}C$ CORD | 800 | 11.0 | 2.0 | 32 | 2800 | 600 | 14.0 | 7.5 | 496.8 | 198.7 | 53.0 | 25 | QSINE | SSB 4 |
| 2D $^{13}C$-DP INADEQUATE | 800 | 6.0 | 2.0 | 16 | 2800 | 680 | 14.0 | 7.4 | 496.8 | 225.8 | | 22 | GM | LB-10, GB0.05 |

**Supplementary Table 3. $^{13}C$ chemical shifts of biomolecules in *S. pombe* cell walls at ambient temperature.** Superscripts are used to denote different allomorphs. Not applicable (/). Branched (Br).

| Carbohydrates | | C1 | C2 | C3 | C4 | C5 | C6 | CO | $CH_3$ | N | Experiment | References |
|---|---|---|---|---|---|---|---|---|---|---|---|---|
| α-1,3-glucan (A) | a | 101.1 | 71.7 | 84.6 | 68.7 | 70.0 | 60.5 | / | / | / | $^{13}C$-$^{13}C$ CORD | Bhanja *et al*. 2014[8] |
| | b | 102.1 | 69.6 | 81.2 | 73.1 | 71.7 | 60.8 | / | / | / | $^{13}C$ DP J-INADEQUATE | |
| | c | 100.2 | 71.1 | 81.3 | 69.6 | 71.8 | 60.9 | / | / | / | | |
| β-1,3-glucan (B) | | 103.6 | 74.4 | 86.8 | 68.3 | 77.5 | 61.3 | / | / | / | $^{13}C$-$^{13}C$ CORD | Shim *et al*. 2007 Fairweather *et al*. 2004 Saito *et al*. 1979[9–11] |
| β-1,3-glucan ($B^{Br}$) | | 103.3 | 73.9 | 85.6 | 69.1 | 76.1 | 69.2 | / | / | / | $^{13}C$ DP J-INADEQUATE | Lowman et al. 2011[12] |
| β-1,6-glucan (H) | | 103.7 | 74.1 | 76.5 | 70.5 | 74.8 | 69.5 | / | / | / | | |
| α-1,2-mannose ($Mn^{1,2}$) | | 101.1 | 80.6 | 71.6 | 67.9 | 74.3 | 61.9 | / | / | / | | Latgé *et al*. 1994 Chakraborty *et al*. 2021[13,14] |
| α-1,6-mannan ($Mn^{1,6}$) | | 101.4 | 72.9 | 73.8 | 67.7 | 72.8 | 66.6 | / | / | / | | |

**Supplementary Table 4. The molar composition of rigid polysaccharides.** The numbers are estimated using integrals (volume) of cross peaks in 2D $^{13}C$-$^{13}C$ 53 ms CORD spectra. The average integrals of cross-peaks of each polysaccharide are shown. Error bars are standard errors.

| **Strains** | **Polysaccharides** β-1,3-glucan (B) | | α-1,3-glucan (A) | | β-1,6-glucan (H) |
|---|---|---|---|---|---|
| | a | b | a | b | |
| wildtype | 64±8 | - | 24±5 | 10±1 | 2±1 |
| *bgs3-1* | 60±7 | 13±5 | 21±5 | 6±1 | - |
| *ghs2-2* | 62±17 | 14±7 | 11±5 | 13±10 | - |
| *bgs3-1 ghs2-2* | 52±11 | 13±2 | 35±16 | - | - |

**Supplementary Table 5. The molar composition of mobile polysaccharides.** The numbers are estimated using integrals (volume) of cross peaks in 2D $^{13}C$-$^{13}C$ refocused DP-J INADEQUATE spectra. The average integrals of cross-peaks of each polysaccharide are shown. Error bars are standard errors of the peak integrals.

| **Strains** | β-1,3-glucan (B) | Branched β-1,3-glucan ($B^{Br}$) | α-1,3-glucan ($A^a$) | α-1,3-glucan ($A^b$) | α-1,3-glucan ($A^c$) | β-1,6-glucan (H) | α-1,4-glucan ($A^{1,4}$) | α-1,2-mannose ($Mn^{1,2}$) | α-1,2-mannan ($Mn^{1,6}$) |
|---|---|---|---|---|---|---|---|---|---|
| wildtype | 4±1 | 12±1 | 19±4 | 4±2 | 11±2 | 20±3 | 2±1 | 17±2 | 11±6 |
| *bgs3-1* | 31±5 | 5±2 | 13±2 | 5±0.2 | 11±2 | 11±1 | 3±0.1 | 13±1 | 8±1 |
| *ghs2-2* | 3±0.3 | 10±4 | 23±2 | 6±2 | 13±2 | 14±5 | 2±1 | 14±4 | 15±4 |
| *bgs3-1 ghs2-2* | - | - | - | - | - | - | - | - | - |
